# The landscape of longevity across phylogeny

**DOI:** 10.1101/2020.03.17.995993

**Authors:** Dmitriy I. Podolskiy, Andrei Avanesov, Alexander Tyshkovskiy, Emily Porter, Michael Petrascheck, Matt Kaeberlein, Vadim N. Gladyshev

## Abstract

Lifespan of model organisms can be extended by genetic, dietary and pharmacological interventions, but these effects may be negated by other factors. To understand robustness of longevity interventions within and across species, we analyzed age-dependent mortality of yeast, fruit flies, nematodes and mice subjected to thousands of genetic, pharmacological or dietary interventions, and applied the principles learned to other organisms. Across phylogeny, the accessible space of lifespan distribution functions, the “landscape of longevity”, has a distinct structure of a fiber bundle, with individual fibers given by Strehler-Mildvan degeneracy manifolds. Within species, most interventions perturb parameters of survival curves along particular degeneracy manifolds. Transitions across manifolds are difficult to achieve, but they may lead to robust lifespan-modulating effects. Analyses of intraspecific degeneracy manifolds revealed soft bounds on achievable longevity. For humans, this bound is ∼138 years.

## Introduction

Studies on the modulation of lifespan of model organisms by genetic, dietary and pharmacological interventions have shown that species longevity, represented by lifespan and the rate of aging, is highly plastic (Kenyon, 2005). It is also known that median and maximal lifespan of experimental cohorts can change significantly in response to interventions; e.g. a null allele of *age-1* in *C. elegans* is characterized by a median lifespan of 145-190 days, ∼10 times more than that of wild-type animals (Ayyadevara et al., 2008), and median lifespan of Ames dwarf mice is extended by ∼50% compared to controls (Bartke et al., 2001a; Brown-Borg et al., 1996). Similarly, the all-cause mortality doubling rate of animals can be increased or decreased by as much as an order of magnitude.

However, the true potential of lifespan-extending interventions is often masked and may be entirely cancelled out by other factors. Examples are clearly seen in large-scale screens, such as the Interventions Testing Program (ITP) of the National Institute on Aging that analyzed the effects of different compounds on mouse lifespan (Warner et al., 2000). While this program reliably detects relatively large effects on longevity, batch effects are evident even across controls, e.g. UM-HET3 males consistently live longer in the animal facilities at University of Michigan than at Jackson Laboratory or University of Texas Health Science Center. Similarly, in the screens aiming to determine the lifespan-extending effects of rapamycin (Harrison et al., 2009; Miller et al., 2011), aspirin (Strong et al., 2008), Protandim (Strong et al., 2016) and nordihydroguaiaretic acid (Strong et al., 2008), strong variability of survival curves was observed across different years study sites. This ambiguity limits applicability of interventions that extend lifespan of animal models to humans.

To address the nature of this variability, we collected survival statistics from large-scale screening studies for interventions affecting lifespan of model organisms, including yeast, nematodes, fruit flies and mice. In addition, we performed our own screening in fruit flies, analyzed publicly available survival data on model organisms and integrated this information with human demographic data. Analyzing the combined dataset, we showed that one fundamental source of the observed variability in aging studies is a high sensitivity of some parameters of survival curves of many species to generic perturbations, combined with a vanishing sensitivity of other parameters to the same perturbations. In other words, it is much easier to perturb some modalities of lifespan distribution functions than others, and even weak interventions, e.g. changes in environmental temperature, can induce such perturbations. Our analysis revealed a global structure of the longevity landscape across interventions affecting lifespan of individual species as well as across different species and uncovered natural soft bounds on the maximal achievable lifespan, which we then quantified for common model organisms and humans. Finally, we performed an extensive information-theoretical analysis, which allowed us to explain the observed structure of the longevity landscape and identify an “extended lifespan” sector, which is characterized by a mortality rate unchanging with age.

## Results

### Defining the structure of longevity landscape across interventions for model organisms

To analyze how lifespan-modulating interventions modify quantitative characteristics of survival and mortality, we first aggregated mortality data for (i) 4,698 strains of *S. cerevisiae* produced by single gene knockouts (McCormick et al., 2015), (ii) 1,416 different single pharmacological compound diets applied to *elegans* (Ye et al., 2014), (iii) 162 single gene knockdowns in *D. melanogaster* that we analyzed for the current study, (iv) ∼15,000 UM-HET3 mice in the Interventions Testing Program (ITP) that subjected animals to diets containing candidate longevity interventions, and (v) various human populations involving different countries and time periods (Human Mortality Database, 2015). Although the survival curves for these 5 species were generally multiparametric complex functions of age, we exploited the well-known observation that the behavior of age-dependent mortality rate for these organisms can be well approximated by the Gompertz function 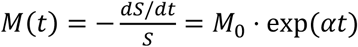 encoding an accelerating decrease of chance of survival with age (Figures 1A-D, Figure 1 – figure supplement 1 - Figure 1 – figure supplement 3). The resulting survival curves were thus parametrized in terms of the initial all-cause mortality rate *M*_0_ and the all-cause mortality doubling rate (“the rate of aging”) *α*, and interventions perturbing these parameters produced lifespan distributions spanning the space (*M*_0_, *α*), further denoted as “the landscape of longevity”.

**Figure 1.**
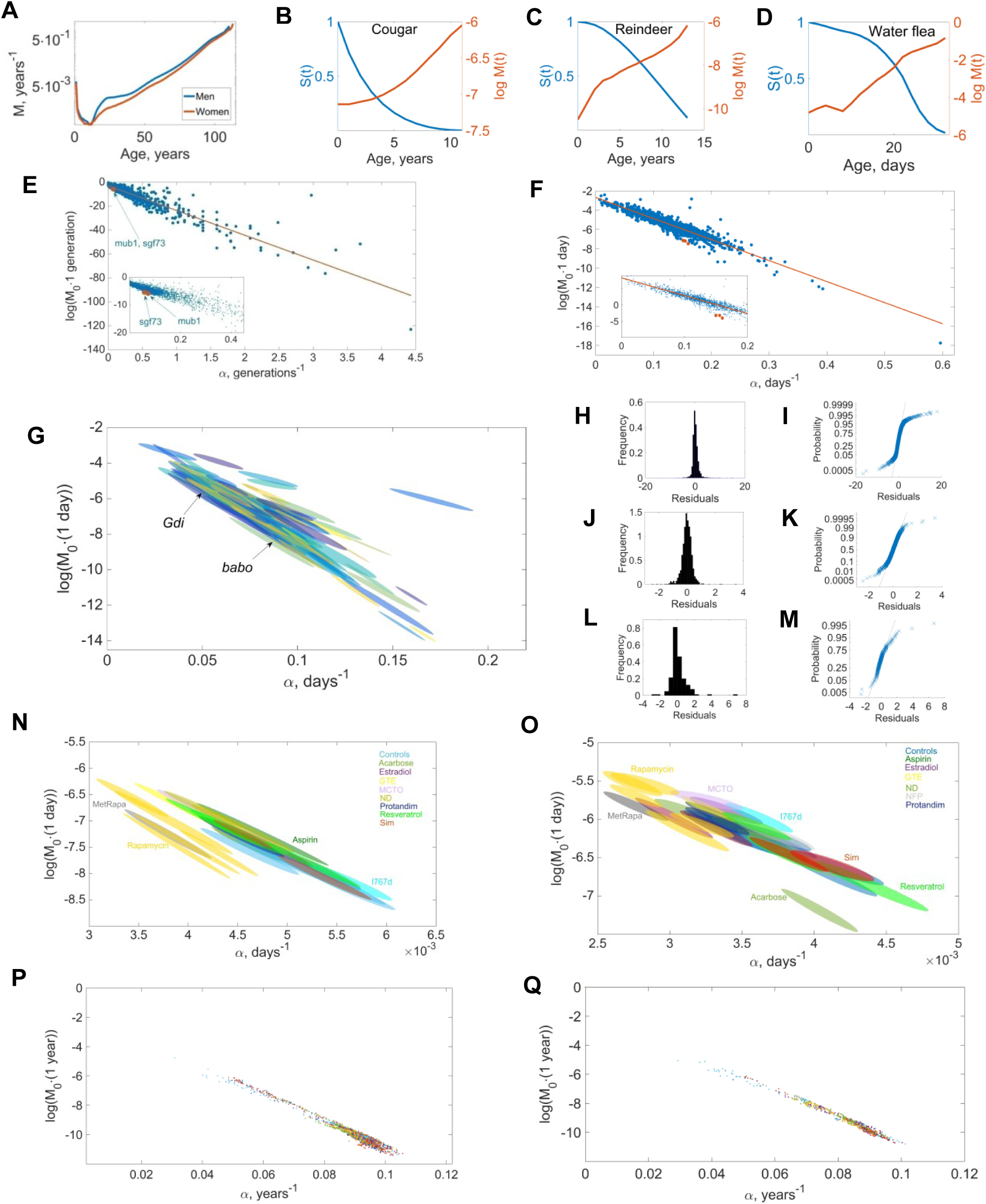
Characteristics of longevity across interventions. **A.** Human mortality rate (men are shown in blue and women in red) grows exponentially with age following the Gompertz law with exceptional precision (*R*^2^ = 0.9602 for women, *R*^2^ = 0.957 for men). As a result, human mortality rates and lifespan distribution functions can be parametrized in terms of initial mortality rate *M*_0_ and the doubling rate *α* (the rate of aging). **B-D.** Gompertz law of mortality accelerating with age also applies to many other organisms, with representative species shown. Age-dependent survival curves *S*(*t*) are shown in blue, and logarithms log *M*(*t*) of the associated mortality rates in red. **E.** Strehler-Mildvan degeneracy manifold on the landscape of parameters (*M*_0_, *α*) of lifespan distribution functions for 4,698 strains of budding yeast, each characterized by a different single gene knockout. (McCormick et al., 2015) Inset shows the behavior of the degeneracy manifold near the origin (region of small *α*). Red dots correspond to two strains (*mub1* and *sgf73* knockouts) with the largest lifespan detected in the screen. **F.** The same for *C. elegans*. Separate points on the plot correspond to parameters of mortality curves for nematodes on different compound diets (1,416 individual diets) (Ye et al., 2014). Inset represents the behavior of the Strehler-Mildvan degeneracy manifold near the origin. Red dots correspond to the diets that resulted in the maximal lifespan observed in the screen. **G.** Degeneracy manifold of survival curve parameters for the lifespan screen of *D. melanogaster*; each dot represents a strain with a specific single gene knockdown. Strains with the maximal lifespans detected in the screen are denoted by arrows (*Gdi* and *babo* knockdowns). **H-I.** In the yeast screen, the vast majority of single gene knockouts led to a displacement of parameters (*M*_0_, *α*) along the degeneracy manifold. Deviations from the degeneracy manifold were approximately Gaussian-distributed (**H**) with a relatively small number of distinct outliers (**I**) characterized by significantly larger or smaller average lifespans. **J-K.** The same for the screen of pharmacological compounds modulating lifespan of *C. elegans*. **L-M.** The same for the screen of single gene knockdowns in the genome of *D. melanogaster* modulating its lifespan. For all three species, interventions modulating lifespan displaced the values of mortality parameters (*M*_0_, *α*) along the degeneracy manifold. **N.** Degeneracy manifold of mortality parameters (*M*_0_, *α*) of ITP mice on diets containing different compounds known to extend their lifespan. Individual ovals correspond to mice on specific diets. Ovals represent errors in identification of the parameters (*M*_0_, *α*). Diets containing rapamycin led to the largest lifespan extension in male mice. **O.** The same for female mice. **P.** Degeneracy manifold of human mortality parameters (men only). Gompertz parameters for men living in different countries are extremely closely distributed along the Strehler-Mildvan degeneracy manifold. Different colored points correspond to actuarian curves for populations of different countries and different decades of 18-21 centuries. **Q.** The same for women.

For all analyzed datasets, species and interventions, we found that the parameters of lifespan distribution were distinctly distributed along particular submanifolds (Figures 1E-G,N-Q), each described by the Strehler-Mildvan correlation line log 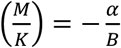 (Lenart and Missov, 2016; Strehler and Mildvan, 1960; Tarkhov et al., 2017a), with *K* and *B* denoting Strehler-Mildvan slope and intercept, respectively. For the screen of 4,698 single gene knockouts affecting yeast replicative lifespan (McCormick et al., 2015) (Figure 1E) we found the Strehler-Mildvan slope of 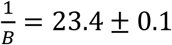 generations and the intercept of log(*K* · 1 *generation*) = −2.761 ± 0.03. The goodness of robust least squares fit for the Strehler-Mildvan model was exceptionally high with *R*^2^ = 0.9781. The goodness of fit of the Strehler-Mildvan model of the Gompertz parameters characterizing survival curves in the pharmacological network screen in *C. elegans* (Ye et al., 2014) was *R*^2^ = 0.9284 (Figure 1F), and the values for the Strehler-Mildvan slope and intercept were 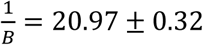 days and log(*K* · 1 *day*) = −2.775 ± 0.045, respectively. We also found that the Gompertz parameters of survival curves for *D. melanogaster* knockdown strains followed the Strehler-Mildvan curve with the goodness-of-fit *R*^2^ = 0.9155 (Figure 1G). Extracting the values of Strehler-Mildvan slope and intercept from the model revealed 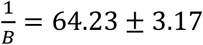 days, log(*K* · 1 *day*) = −1.759 ± 0.278.

Survival data of ITP male mice on diets containing different pharmacological compounds (Figure 1N) revealed that the Gompertz parameters followed the Strehler-Mildvan correlation with g.o.f. *R*^2^ = 0.5221, and the values of the Strehler-Mildvan slope and intercept were 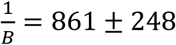 days, log(*K* · 1 *day*) = −3.853 ± 0.877. For female mice (Figure 1O), we found the 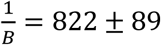 days, log(*K* · 1 *day*) = −4.242 ± 0.397, the g.o.f. *R*^2^ = 0.9672 for the Strehler-Mildvan correlation. Finally, the human data similarly followed the Strehler-Mildvan law with g.o.f. *R*^2^ = 0.97 for both men (Figure 1P) and women (Figure 1Q). For men, we found log(*K* · (1 *year*)) = −1.512 ± 0.009, 1/*B* · (1 *year*)^−1^ = 92.38 ± 0.11, and for women the Strehler-Mildvan slope and intercept were given by log(*K* · (1 *year*)) = −1.396 ± 0.069 and 1/*B* · (1 *year*)^−1^ = 95.58 ± 0.78. Combining mortality data for men and women, the resulting Strehler-Mildvan slope and intercept were found to be log(*K* · (1 *year*)) = −1.345 ± 0.031 and 1/*B* · (1 *year*)^−1^ = 95.49 ± 0.36.

Importantly, for all considered model organisms the vast majority of pharmacological interventions, gene knockouts/knockdowns and diets induced displacements of parameters of survival curves along the Strehler-Mildvan degeneracy manifolds corresponding to a given species, while displacements away from them were Gaussian-distributed and relatively small (Figures 1H-M). However, there were also rare outliers (Figures 1I,K,M) typically corresponding to strong displacements away from the Strehler-Mildvan degeneracy manifold. For humans, individual points in the (*M*_0_, *α*) space of lifespan distribution parameters were determined using demographic data, and variance in the values of (*M*_0_, *α*) for different points was probably due to different lifestyle, climate, diet, time periods, etc. Nevertheless, the same pattern of Strehler-Mildvan degeneracy was observed for humans as for other analyzed species. Thus, the majority of interventions modulating lifespans and the rates of aging in model animals and humans lead to a displacement of lifespan distribution parameters along the Strehler-Mildvan degeneracy manifolds, although some interventions may lead to transitions between different degeneracy manifolds.

### Defining the structure of longevity landscape across species

To characterize the relation between degeneracy manifolds in the global longevity landscape of parameters (*M*_0_, *α*), we further aggregated publicly available data on lifespan-extending interventions in *C. elegans, melanogaster, M. musculus* and *R. norvegicus* (Supplementary File 2). Although the data were collected from different sources, for all considered species the parameters of survival curves turned out to be again well fitted by the Strehler-Mildvan correlation lines within individual species (Figure 2A). We found that (i) survival curves for individual species are typically characterized by existing degeneracy manifolds, and the largely continuous transition between the points on the longevity landscape corresponding to different species corresponds to the transition between different degeneracy manifolds; (ii) longer-lived species are characterized by the degeneracy manifolds with higher slopes; and (iii) for shorter-lived species, the observed relative deviations from their corresponding Strehler-Mildvan manifolds were much larger than for longer-lived species.

**Figure 2.**
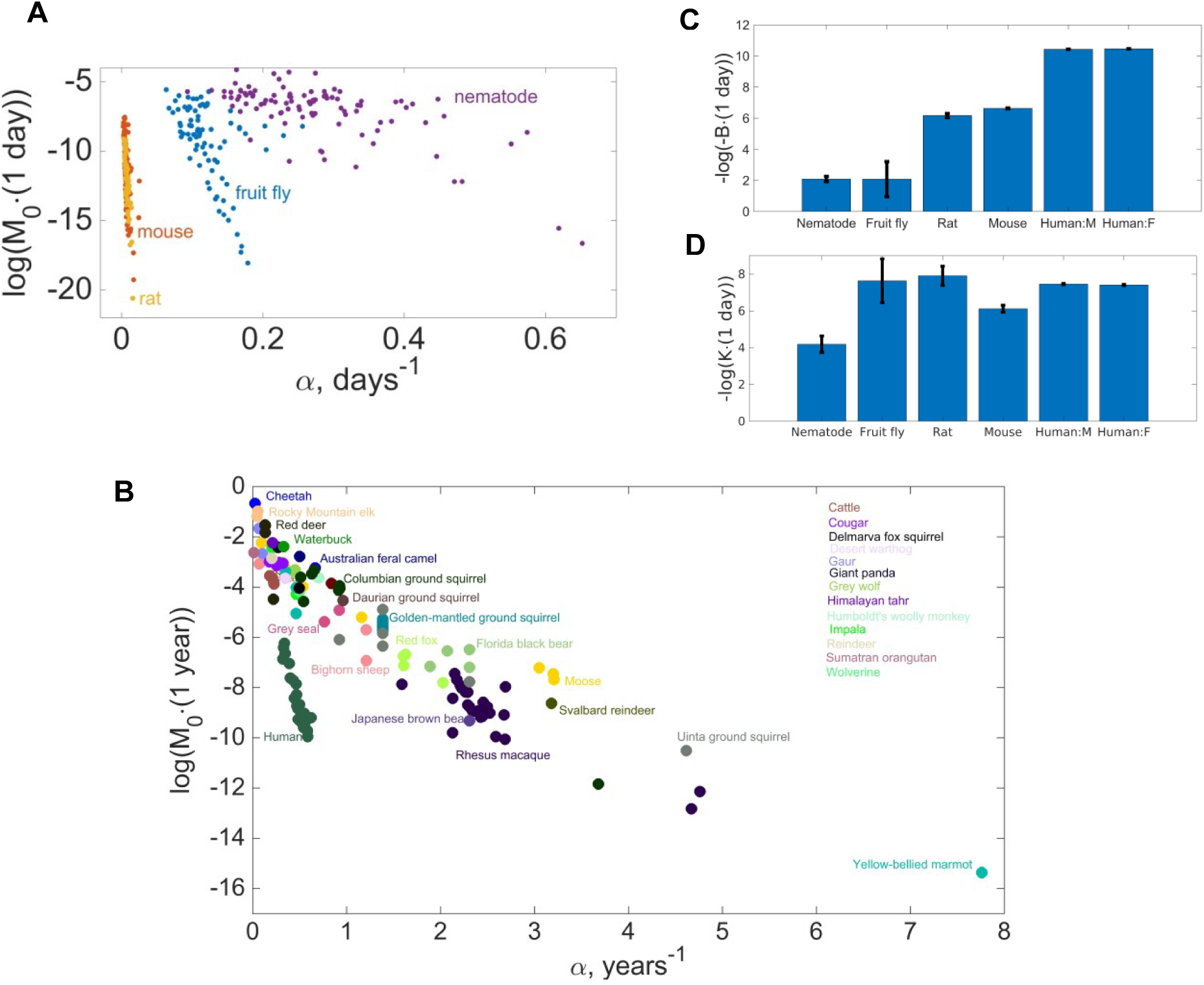
Characteristics of longevity across species. **A.** Individual degeneracy manifolds and their colocation for mice, rats, fruit flies and nematodes. Each point represents mortality rate parameters (*M*_0_, *α*) from a separate experiment involving testing a particular intervention modulating a species lifespan. Data collected from literature (Table S2). **B.** Structure of the landscape of mortality rate parameters across 90 species (longer-lived species are shown). Note that the value of the slope of the degeneracy manifold for humans is significantly higher than for other species. **C.** Strehler-Mildvan parameter *B* (slope of the degeneracy manifold in the space of parameters (*M*_0_, *α*)) for nematodes, fruit flies, mice, rats and human (men and women, separately). The value of *B* depends strongly on the average lifespan of the species. **D.** The same for the Strehler-Mildvan parameter *K* (intercept of the degeneracy manifold in the space of parameters (*M*_0_, *α*)). Unlike the slope *B*, the value of intercept *K* remains nearly constant for different species, implying that Strehler-Mildvan manifolds are parametrized by a single parameter *B* rather than two independent parameters *K* and *B*.

To further examine the structure cross-species longevity landscape, we combined the survival data for 90 relatively long-lived species (including many mammals) with the age-dependent behavior of mortality approximately following the Gompertz law (Salguero-Gómez et al., 2016) (Figure 2B). For many species, Gompertz parameters of survival curves were clustered close to a single Strehler-Mildvan-like degeneracy manifold, representing a global attractor on the longevity landscape across species. This demonstrated that even very significant perturbations of the genome leading to the transition across species still may correspond to the displacement of longevity parameters (*M*_0_, *α*) along the same degeneracy manifold.

### Strehler-Mildvan degeneracy manifolds are one-parametric

Performing quantitative analysis of the structure of the longevity landscape (Figure 2B), we compared the values of the Strehler-Mildvan slope and intercept across analyzed species. This analysis revealed that (i) the slope *B* values differed significantly across species, with longer-lived species characterized by larger values of inverse Strehler-Mildvan slope 1/*B* (Figure 2C; note that the scale is logarithmic, so the difference in 1/*B* between species is exponentially large in the linear scale), while (ii) the values of the Strehler-Mildvan intercept *K* remained relatively constant. This finding suggests that the structure of Strehler-Mildvan degeneracy manifolds on the longevity landscape is approximated by one-parametric (defined by a single free parameter *B* discriminating individual degeneracy manifolds) rather than two-parametric (defined by the Strehler-Mildvan parameters *K* and *B*) functions. The structure of longevity landscape across species (Figure 2B) also supported this conclusion, as points corresponding to particular long-lived species (e.g. human, rhesus macaque, black bear, red fox, etc.) were visually combined into different Strehler-Mildvan degeneracy manifolds converging towards the same point/area at the origin of the plot corresponding to the vanishing all-cause mortality doubling rate *α* = 0.

### Achievable maximal lifespan is bounded for translations along a degeneracy manifold

We further studied the behavior of maximal lifespan achieved in the screens for longevity interventions in yeast, nematodes and fruit flies (in what follows, such lifespans are denoted as *LS*_*max*_(*α*)). In all analyzed screens, maximal lifespans *LS*_*max*_(*α*) obtained in individual experiments appeared to be capped regardless of lifespan-modulating perturbation (Figures 3A-C). In particular, the maximal replicative lifespan of yeast across different interventions in the screen of 4,698 single gene knockouts was capped at 87 generations (Figure 3A), maximal lifespan of nematodes in the pharmacological screen at 48 days (Figure 3B) and maximal lifespan of *D. melanogaster* in the screen of 162 gene knockdowns at 105 days (Figure 3C). In each case, we observed that maximal achieved lifespans *LS*_*max*_(*α*) remained generally small in experiments leading to cohorts with large mortality doubling rates *α* and start to grow when *α* decreases. However, the value of *LS*_*max*_(*α*) typically reached a saturating bound at a small *α* = *α*_*min*_, and kept decreasing further with decreased *α*. In other words, the lowest *α* values were characterized by lower, not higher lifespan compared to more intermediate values.

**Figure 3.**
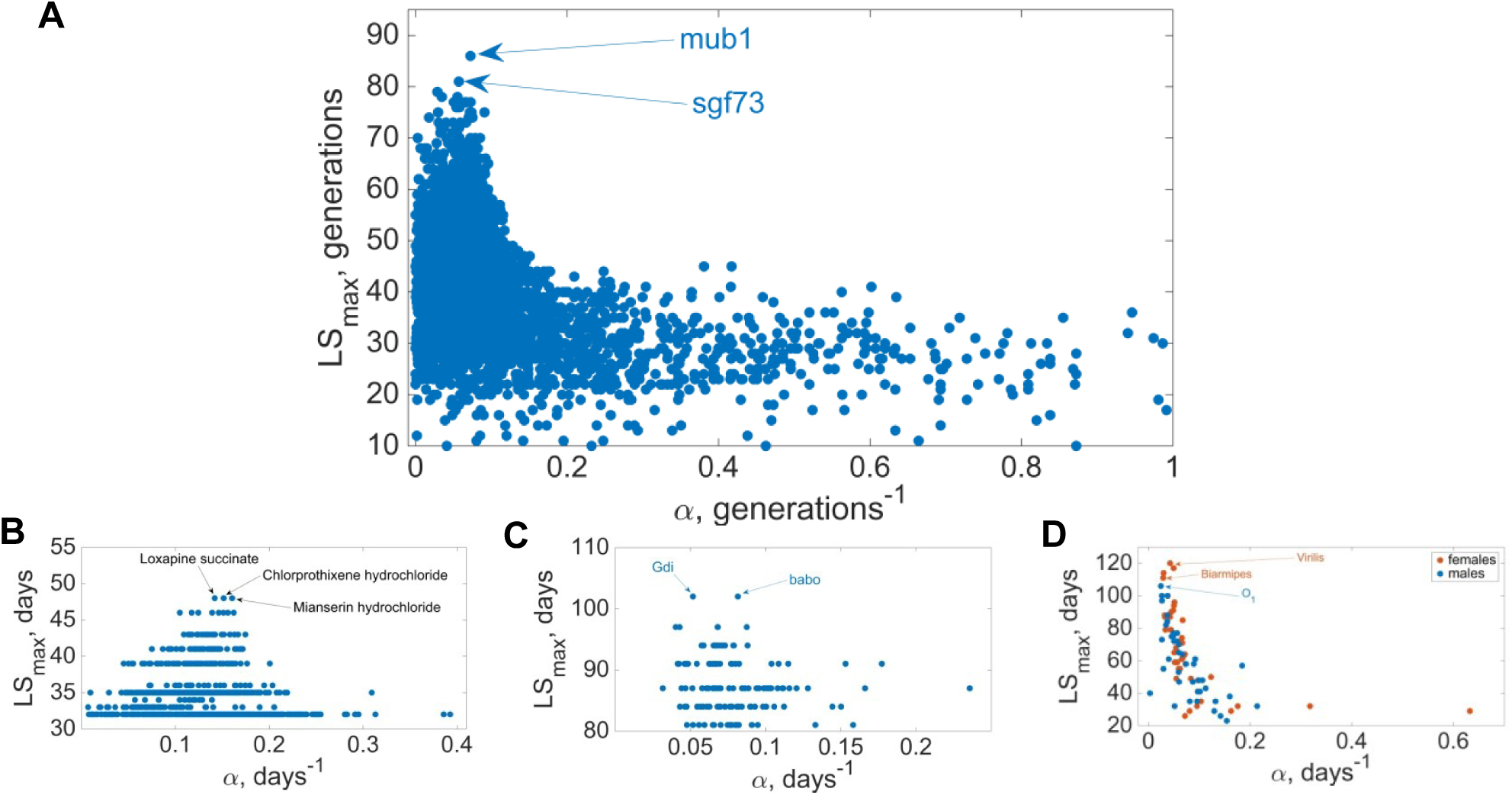
Limits on achievable maximal lifespan across interventions and species. The range of maximal lifespans observed in longevity screens involving genetic and environmental perturbations shows a distinct maximum when plotted as a function of all-cause mortality doubling rate *α*. **A.** Maximal replicative lifespans *LS*_*max*_(*α*) for 4,685 yeast single-gene knockouts. The orange circle denotes the estimate for the achievable replicative lifespan of *S. cerevisiae* along the degeneracy manifold. The maximal lifespan achieved in the screen is 80-95 generations (*mub1* and *cgf73* knockouts). **B.** Maximal lifespan *LS*_*max*_(*α*) of *C. elegans* subjected to 1,416 diets containing individual compounds. In the screen, the observed maximal lifespan was 48 days (*C. elegans* on loxapine succinate, chlorprothixene hydrochloride and mianserin hydrochloride diets). **C.** Maximal lifespan *LS*_*max*_(*α*) for 162 single gene knockdowns of *D. melanogaster*. The maximal lifespan observed in the screen (108 days) was for *babo* and *Gdi* knockdowns. **D.** Dependence of the maximal lifespan *LS*_*max*_ on the value of mortality doubling rate *α* for 13 different species of flies. The structure of the *LS*_*max*_(*α*) across different species of *Drosophila* is the same as in Figures 3A-C: there is a distinct maximum at a certain relatively small value of *α*, and the value of the maximal lifespan drops quickly away from this value. Blue dots correspond to male, and red to female flies.

Similarly, we found that the same applies to maximal lifespans *LS*_*max*_(*α*) for variations between species close to each other in the sense of phylogenetic distance. For example, this was the case for 14 *Drosophila* species (Figure 4D), with *LS*_*max*_(*α*) rapidly increasing with decreasing aging rate *α*, reaching a global maximum at max(*LS*_*max*_) ≈ 110 days for *D. virilis* and rapidly decreasing again with the decreasing *α*.

**Figure 4.**
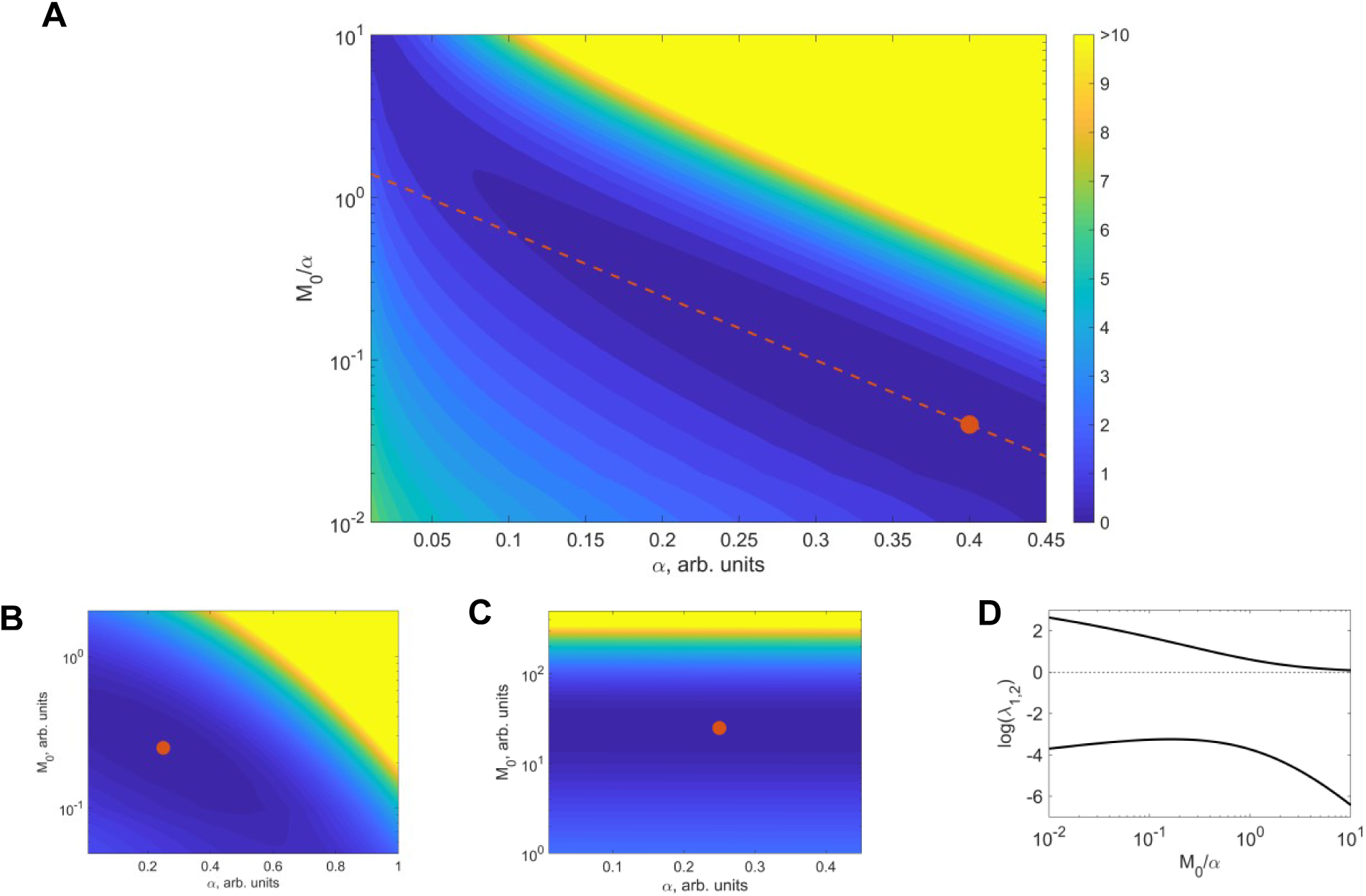
Information-theoretical analysis of survival curves. **A.** Kullback-Leibler (KL) divergence between the control lifespan distribution function (denoted by the red blob; corresponding to *M*_0_ = 0.016, *α* = 0.4, the case 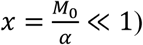) and lifespan distribution functions corresponding to different values of initial mortality *M*_0_ and mortality doubling rate *α*. KL divergence between two distribution functions is an information-theoretical measure of their similarity (Supplemental Information, Section 1). Different colors correspond to different values of KL divergence between a given lifespan distribution and the control distribution; darker colors correspond to distributions more similar to the control distribution. The red line corresponds to the theoretical degeneracy line (3), a one-parametric representation of the Strehler-Mildvan correlation. **B.** Behavior of KL divergence in the case 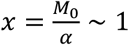. The one parametric Strehler-Mildvan line no longer represents the form of degeneracy manifold well. The degeneracy line in the space of parameters (*M*_0_, *α*) now connects the control distribution and the distribution corresponding to constant mortality rate *α* = 0. **C.** Behavior of KL divergence in the case 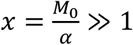. Degeneracy of survival curves in the space of Gompertz parameters *M*_0_ and *α* is now well represented by the line *M*_0_ = *Const*. **D.** Eigenvalues of the Fisher information matrix relating two Gompertz distributions never become of the same order implying existence of a degeneracy manifold at arbitrary values of *x* = *M*_0_/*α* (along the eigenvector of the Fisher information matrix corresponding to its lowest eigenvalue among the two).

### Kullback-Leibler divergence of lifespan distributions

We sought to develop a single theoretical framework to explain and unify the observations outlined above, specifically the findings that (i) most interventions modulating aging in model organisms lead to a displacement of parameters (*M*_0_, *α*) of lifespan distributions along particular Strehler-Mildvan degeneracy manifolds on the landscape of longevity, (ii) Strehler-Mildvan degeneracy manifolds are one-parametric, with different manifolds converging to each other in the limit of slow aging rates *α* → 0, and (iii) maximal lifespan achievable by displacements along particular Strehler-Mildvan degeneracy manifolds is capped.

To develop such a framework, we performed information-theoretical analysis of Gompertz lifespan distribution functions across several species as the all-cause mortality data used in our study fit the Gompertz law of accelerated mortality exceptionally well with *p*-values for the linear fit of logarithm of all-cause mortality to age below 10^−10^ for humans and below 10^−3^ for other species (Figures 1A-D,Figure 1 - figure supplement 1 – Figure 1 – figure supplement 4). This analysis is not very sensitive to a particular functional form of lifespan distribution (Supplementary File 1, Section 9), allowing weak deviations from the Gompertz law such as slowdown of mortality acceleration with age, which are indeed observed for many species.(Jones et al., 2014) However, what is crucial for the conclusions of this and the next sections is the fact that all-cause mortality rate is accelerated with age (also consistent with the behavior of many molecular markers of aging (Petkovich et al., 2017; Podolskiy et al., 2016)). As explained below in “Non-Strehler-Mildvan regime on the landscape of longevity”, species with survival characterized by constant, age-independent mortality rates can also be analyzed within the developed framework.

The lifespan distribution function corresponding to Gompertz-like mortality rate and normalized for very large cohorts is given by the expression 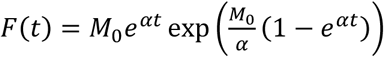 (Figure 1 – figure supplement 5). Any comparative survival analysis is essentially based on identifying deviations in survival function *F*(*t*) between a control cohort and cohorts subjected to interventions modulating control lifespan and the rate of aging. Across all possible comparison measures between two lifespan distribution functions, only the Kullback-Leibler divergence (also known as relative entropy of two distributions or information gain (Kullback and Leibler, 1951)) satisfies canonical properties of entropy and mutual information (Hobson, 1987) (Supplementary File 1, Section 1). Thus, we based our information-theoretical analysis on evaluating the Kullback-Leibler divergence between two lifespan distributions.

We were able to obtain the expression for Kullback-Leibler divergence 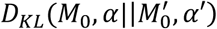 of Gompertz lifespan distribution functions in a closed analytic form reading

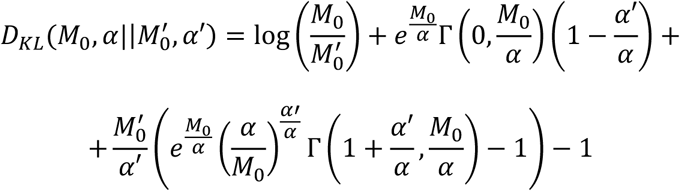

(see also Supplementary File 1, Section 1). Its structure revealed an important caveat in the process of fitting of observed lifespan distribution functions to survival curves *F*(*t*): in the space of fitting Gompertz parameters 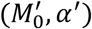 there exist extended regions encompassing distributions, which do not deviate from the control distribution (*M*_0_, *α*) in terms of the Kullback-Leibler divergence *D*_*KL*_ (Figures 4A-C).

The origin of these regions can be revealed by considering Kullback-Leibler divergence of weak deviations from the control lifespan distribution. Namely, it is well known that one can write 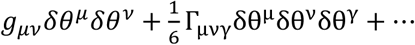, where *g*_*μv*_ is the Fisher information matrix of Gompertz lifespan distribution functions, and *δθ* = (*δM*_0_, *δα*) denotes deviations in the variable parameters 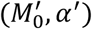 of the Gompertz lifespan distribution *F*(*t*) from the control values (*M*_0_, *α*). As we were able to obtain the Fisher information matrix for Gompertz distribution in the closed form as well (Supplementary File 1, Section 1), we explicitly found that one of the eigenvalues of the Fisher information matrix *g*_*μv*_ remains much smaller than the other one (Figure 4C) for an arbitrary choice of parameters *M*_0_ and *α* (in particular, for any value of the dimensionless parameter *x* = *M*_0_/*α* controlling behavior of the eigenvalues and eigenvectors of the Fisher information matrix). In particular, we found in the case *x* ≪ 1 (realized for virtually all species and interventions analyzed in this study) that a smaller eigenvalue of the Fisher information matrix vanishes logarithmically as 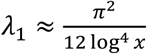, while the larger one grows logarithmically as *λ*_2_ ≈ log^2^ *x* (Supplementary File 1, Section 1). The eigenvector corresponding to a larger eigenvalue of the Fisher information matrix then establishes a direction in the space of parameters (*M*_0_, *α*) such that all Gompertz lifespan distributions obtained by translation along this direction coincide with the control distribution in the sense of Kullback-Leibler divergence between them (the latter vanishes to the leading order in small *δα* and *δM*_0_).

The directions of degeneracy on the fitting landscapes of models corresponding to vanishing eigenvalues of Fisher information matrices are known as “sloppy” (Machta et al., 2013). This “sloppiness”, i.e. sensitivity of distributions to perturbations of their parameters, implies that even relatively weak perturbations of the data fitted to the distributions with “sloppy” directions on the fitting landscape lead to strong deviations of the resulting fit along the directions of degeneracy. Similarly, an intervention physically affecting the actual biological system behind the obtained distribution (lifespan distribution in our case) would easily induce a perturbation along the degeneracy manifolds on the landscape of parameters (*M*_0_, *α*), while the perturbation across different degeneracy manifolds would be much harder to achieve. This explains the fact that most lifespan-modulating interventions analyzed in this study led to displacements of lifespan distribution function parameters (*M*_0_, *α*) along particular degeneracy manifolds.

Physical interventions of interest would naturally include arbitrary systematic changes in conditions under which the analyzed cohort of model animals is held: temperature, variation of diet or handling conditions, etc. All such effects would thus again lead to perturbations in the characteristics of cohort survival, to be translated into large perturbations in the space of parameters (*M*_0_, *α*) along the degeneracy manifolds. To reiterate, as sensitivity of lifespan distributions with respect to perturbations along such “sloppy” directions on the longevity landscape is extremely high, it is easy to achieve changes in lifespan distributions associated with such perturbations, even when such perturbations are weakly changing the biological markers of the organisms in the cohort. Such weak perturbations would, for instance, include location of the lab where analysis is performed or minor differences in the reagents used. In our opinion, this effect is one of the major sources of variability in aging studies.

Finally, we note that if the main effect of a particular intervention is a displacement of the lifespan distribution along the degeneracy manifold on the longevity landscape, it should also be relatively easy to introduce a counteracting effect (such as change in environment) leading to the displacement of parameters (*M*_0_, *α*) back to the values of 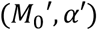 corresponding to the control distribution. On the other hand, interventions leading to displacements of parameters (*M*_0_, *α*) between different degeneracy manifolds are expected to lead to a robust lifespan-extending effect.

### Identifying the physical meaning of Strehler-Mildvan correlation

By fixing a control value of Gompertz parameters (*M*_0_, *α*) and deriving the functional form of the degeneracy manifold corresponding to lifespan distributions located within zero Kullback-Leibler divergence from the control distribution (Supplementary File 1, Section 1), we found that the latter is reduced to

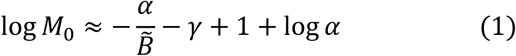

in the limit 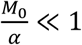, where *γ* = 0.57721566 … is the Euler-Mascheroni constant, and 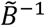 is the parameter with dimension of time enumerating degeneracy manifolds on the landscape of control parameters (*M*_0_, *α*). The functional dependence of *M*_0_ (initial mortality) on *α* (aging rate) is exceptionally close to the Strehler-Mildvan (SM) dependence 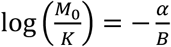 (Strehler and Mildvan, 1960), although it is one-parametric unlike the latter. Thus, we can explain explicitly why Strehler-Mildvan manifolds corresponding to different species as diverse as nematodes, fruit flies, mice and humans, are characterized by close values of the Strehler-Mildvan intercept *K* (Figures 2A-D).

From the expression (1), it becomes clear that the physical meaning of the Strehler-Mildvan slope *B* is the index of different degeneracy manifolds on the fitting landscape of lifespan distribution functions. On the other hand, the Strehler-Mildvan intercept *K* remains *α*-dependent. This should be naturally expected, since if the Strehler-Mildvan correlation is a degeneracy artifact (Burger and Missov, 2016; Tarkhov et al., 2017b), all free parameters entering this correlation should themselves depend on the values of *M*_0_ and *α* only.

### Degeneracy manifolds and temporal scaling

It was previously observed for *C. elegans* that temperature variation and various genetic perturbations lead to scaling deformation of the hazard functions *M*(*t*) (Stroustrup et al., 2016). In particular, for two different environmental temperatures *T*_0_ and *T*_1_ a relationship of the form 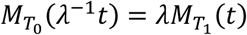 is expected to take place, where *λ* is a dimensionless scale factor. As temperature change is a common environmental variation, its major effect on mortality is the displacement of parameters (*M*_0_, *α*) of survival distribution function along a particular Strehler-Mildvan degeneracy manifold (1). Then, if temperature variation as a weak perturbation leads to MRDT scaling *α* → *λα*, it follows from the expression (3) that the corresponding modification of the hazard function gives the scaling transformation (4) and a temporal shift 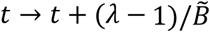, where 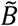 is the constant numerating different degeneracy manifolds (Supplementary File 1, Section 4). Therefore, the phenomenon of temporal scaling of survival curves (Stroustrup et al., 2016) can be explained by the existence of degeneracy manifolds in the space of survival functions and sensitivity of survival functions with respect to generic perturbations across these manifolds.

### Estimating maximal lifespan for perturbations along individual degeneracy manifolds on the longevity landscape

Next, we proceeded to analyze the behavior of maximal lifespan achievable by translations of parameters (*M*_0_, *α*) of survival curves along particular degeneracy manifolds using the obtained expressions for Kullback-Leibler divergence between Gompertz lifespan distribution functions and the corresponding Strehler-Mildvan degeneracy manifolds (Supplementary File 1, Section 7). We found that for translations along such manifolds both average and maximal lifespans grow monotonically with decreasing *α*. However, *α* cannot be generally decreased to 0 along the degeneracy manifold, as the Kullback-Leibler divergence between the realized lifespan distribution function and that of control distribution starts to grow (Figure 4A). Thus, for every degeneracy manifold/Strehler-Mildvan correlation line with 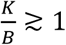 on the landscape of longevity, a value of *α* = *α*_*min*_ exists such that the condition of “sloppiness” for the given degeneracy breaks down, and it becomes hard to push the value of aging rate *α* below the value of *α*_*min*_; this minimal value given by

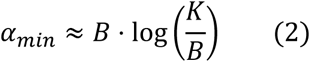

determines the cap on the largest average and maximal lifespan values achievable by perturbations displacing survival distributions along the given degeneracy manifold characterized by the Strehler-Mildvan slope *B* and intercept *K*. Namely, we found (Supplementary File 1, Section 7) that

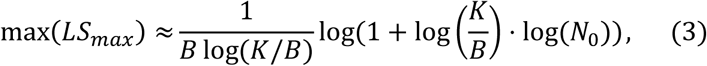

where *N*_0_ is size of the initial cohort (as dependence of the maximal achievable lifespan on *N*_0_ is double-logarithmically weak, the estimate (3) is robust with respect to even significant changes in the size of the cohort). We then proceeded to estimate the value of the cap (3) for several model organisms and humans. For all estimations provided below, the value *N*_0_ = 10^12^ for the initial cohort was taken, and all results were only very weakly sensitive to the value of *N*_0_.

Using parameters *K* and *B* of the Strehler-Mildvan fit for the replicative lifespan of *S. cerevisiae*, we found that the maximal lifespan bound for interventions moving parameters of lifespan curves along the Strehler-Mildvan degeneracy manifold represented on Figure 1E is max(*LS*_*max*_) ≈ 147 generations.

For the pharmacological screen of *C. elegans*, we found that the corresponding bound is max(*LS*_*max*_) ≈ 167 days. This significantly exceeds the maximal lifespan achieved in the screen (48 days for the diets containing mianserin, chlorprothixene and loxapine), but is also lower than the lifespan of *age-1* null mutants (Ayyadevara et al., 2008) (∼230 days). Also, cohorts with the maximal lifespan of 72 and 75 days were detected in a similar larger-scale lifespan screen testing ∼88,000 small molecule diets and over 1 million animals (Petrascheck et al., 2007). However, a careful analysis shows that the parameters of mortality curves for 6x outcrossed *age-1* mutants (the strain characterized by the 10-fold lifespan extension) are located away from the main Strehler-Mildvan degeneracy manifold/fiber (Figure 2B). Indeed, the Gompertz fit of the mortality data from (Ayyadevara et al., 2008) reveals *α* = 0.03968 ± 0.02306 days and log(*M*_0_ · (1 *day*)) = −6.164 ± 1.98 (g.o.f. *R*^2^ = 0.566), corresponding to a point on the Strehler-Mildvan manifold significantly below the Strehler-Mildvan degeneracy manifold presented in Figure 1F. This indicates that the *age-1* null mutation leads to a strong robust perturbation of survival curves allowing to leave the main Strehler-Mildvan degeneracy manifold of *C. elegans*. It would be interesting to determine whether other alleles of *age-1* similarly fall off the main degeneracy line for *C. elegans* on the longevity landscape. As was mentioned above, we also collected mortality data from many previous experiments on lifespan modulation in *C. elegans* (Supplementary File 2, Figure 2 – figure supplement 1). Their analysis revealed that, for most experiments described in the literature, parameters of the corresponding mortality curves belong to the Strehler-Mildvan degeneracy manifold identified above (Figure 2A).

For the genetic screen of 162 single gene knockdowns in *D. melanogaster*, we found that the value of maximal lifespan achievable along the degeneracy manifold is max(*LS*_*max*_) ≈ 112 days. In our screen, the age of 110 days which is close to this bound was reached only in three cohorts (*Gdi, babo* and *CG11877* knockdowns). It is interesting to compare these numbers to those achieved in the screen of 14 *Drosophila* strains and species, including *melanogaster* (*yw* strain and three longevity-selected strains *B3, O1* and *O3*), *biarmipes, erecta, mojarensis, pseudoobscura, saltans, sechelia, simulans, virilis, willistoni* and *yakuba* as well as the same lines subjected to two forms of calorie restriction (Figure 4D). Gompertz fit parameters for survival curves of the strains followed the Strehler-Mildvan correlation with the goodness-of-fit *R*^2^ = 0.6702 and the values of the Strehler-Mildvan slope and intercept given by 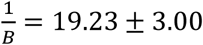 days, log(*K* · 1 *day*) = −2.87 ± 0.34. The maximal achievable lifespan was thus max(*LS*_*max*_) ≈ 271 days. Among the species included in the screen, only the long-lived *Drosophila virilis* strain in fact reached the age of 123 days. The difference between estimations of maximal lifespan achievable along the main degeneracy manifold for *D. melanogaster* and in a multi-species screen may be related to the fact that laboratory experiments on *D. melanogaster* use species evolved for hundreds of generations in laboratory conditions favoring early reproduction and thus lived significantly shorter that wild-caught *D. melanogaster*. We also analyzed many studies on lifespan modulation that used genetic interactions (Supplementary File 2, Figure 2 – figure supplement 2). While parameters of mortality curves were found to belong to the same SM correlation line/fiber identified above, the mortality curves of a different *D. melanogaster* strain subjected to knockout of genes from the *Tor* complex seemed to form a separate SM fiber corresponding to slightly extended lifespans (Figure 2 – figure supplement 2).

For male mice (Figure 1N), we found that the maximal lifespan achievable by translations along the corresponding Strehler-Mildvan degeneracy manifold was given by max(*LS*_*max*_) ≈ 1303 days (∼ 3.6 years). For females (Figure 1O), on the other hand, it was max(*LS*_*max*_) ≈ 1476 days (∼ 4.01 years). It is interesting that the max(*LS*_*max*_) is close to the lifespan achieved in Ames mice (Brown-Borg et al., 1996), although the bound reported here is not very accurate. Previously, it was demonstrated (Bartke et al., 2001b) that calorie restriction extends the lifespan of Ames mice to ∼1,400 days, which is also close to the bound reported here.

We further combined these results with the mortality curves from various experiments on lifespan modulation in mice (Supplementary File 2, Figure 2 – figure supplement 3, Figure 2 – figure supplement 4). Again, parameters of mortality curves produced by the studies involving different gene knockouts belonged to the main Strehler-Mildvan degeneracy manifold, and for various calorie and dietary restriction experiments such parameters were located significantly below the main manifold. We also noted that among different interventions calorie restriction stands apart, with the parameters of survival curves forming a distinctly different degeneracy manifold on the landscape of longevity.

Finally, using the data from the Human Mortality Database and the obtained parameters of the Strehler-Mildvan degeneracy manifold for humans (Figures 1P,Q) we found that the maximal lifespan achievable by translations along this manifold (men and women combined) was max(*LS*_*max*_) ≈ 138 years. This result is ∼20% above that recently estimated (Dong et al., 2016a) and ∼16% above the maximum lifespan observed. It should be noted that while previous analyses (Dong et al., 2016a) focused on extreme long-lived outliers, we consider fundamental features of mortality patterns(as well as survival curves describing mortality rates in large cohorts including individuals of different ages) to arrive at the conclusion.

We would like to emphasize that while the analysis presented here is fundamentally based on the Gompertz mortality law, the statement about the existence of degeneracy manifolds in the space of parameters *α* (mortality rate) and *M*_0_ (initial mortality) holds true generally for mortality curves and lifespan distributions significantly deviating from Gompertz-like but preserving the mortality rate acceleration with age. However, it is also well known that, for many species, exponential growth of mortality rate *M*(*t*) slows down at advanced ages, and *M*(*t*) reaches plateau at very late ages (Carey et al., 1992; Vaupel et al., 1998) (recently, it was also argued for humans (Barbi et al., 2018)). This regime of plateau requires a separate and involved analysis which will be done elsewhere.

### Non-Strehler-Mildvan regime on the landscape of longevity

As was mentioned above, initial mortality rate *M*_0_ and aging rate*α* characterizing lifespan distribution functions for all species and virtually all interventions modulating their lifespan considered in this study are such that the dimensionless ratio of *M*_0_ and *α* is very small: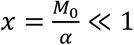. Analysis of the explicit expression for the Kullback-Leibler divergence between two Gompertz lifespan distributions (Supplementary File 1, Sections 1-3) shows that for every such pair of parameters (*M*_0_′, *α*′) a Strehler-Mildvan-like degeneracy manifold exists in the space of possible (*M*_0_, *α*) such that the vast majority of interventions perturbing lifespan of the considered species leads to displacement (*δM*_0_, *δα*) from the point (*M*_0_′, *α*′) along this manifold. As *x* grows, the Kullback-Leibler divergence *δD*_*KL*_ between the distributions (*M*_0_, *α*) and (*M*_0_′, *α*′) starts to grow as well (Figure 4B), and *δD*_*KL*_ becomes of the order 1 at a certain *x* inducing the bound on the maximal lifespan discussed in the previous Section. This growth is due to the presence of terms of higher order than quadratic in (*δM*_0_, *δα*) in the Kullback-Leibler divergence *δD*_*KL*_, which become important at *x* ≳ 1 (Supplementary File 1, Sections 2 and 7). The majority of generic perturbations (*δM*_0_, *δα*) lead to a displacement of parameters (*M*_0_, *α*) along the degeneracy manifold (1) with a bias for displacements towards larger *α* (as the regime of small *α* is hard to reach due to the growth of *δD*_*KL*_).

One may inquire what happens if the control distribution (*M*_0_′, *α*′) is such that 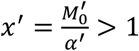. We found that the behavior of perturbations (*δM*_0_, *δα*) in this regime is distinctly different from that described above (Figure 4C, Supplementary File 1, Section 5). The degeneracy manifold on the landscape of parameters (*M*_0_, *α*) is now determined by the condition *M*_0_ = *Const*; it is unbounded at small *α* (in the sense that *δD*_*KL*_ does not grow along the degeneracy manifold even as *α* → 0) but becomes bounded by the condition *δD*_*KL*_ ∼ 1 at large *α* or *x* ∼ 1. Therefore, perturbations (*δM*_0_, *δα*) will generally lead to a displacement of parameters (*M*_0_, *α*) along the line *M*_0_ = *Const* easily reaching the limit (*M*_0_, *α* = 0) corresponding to mortality rate remaining constant with age, i.e., the regime of mortality plateau,(Barbi et al., 2018; Carey et al., 1992) although the points (*M*_0_, *α*) with *α* ≪ *M*_0_ corresponding to a slow Gompertz-like growth of mortality rate with age will be also accessible.

Existence of this regime on the landscape of longevity parameters across species is an unavoidable consequence of the theory explaining the observed structure of the longevity landscape. While we have not observed examples of such regime realized in the experiments analyzed here, possible cases may include species with mortality rates featuring plateau or species with very low rates of aging (Jones et al., 2014) such as the naked mole rat (Buffenstein, 2008; Kim et al., 2011; Ruby et al., 2018). As described above, parameters of survival and mortality of such species would be characterized by exceptional stability under general perturbations including the effects of lifespan-modulating genetic, pharmacological and dietary perturbations, which is a testable prediction of the theory outlined above.

## Discussion

We found that topology of the longevity landscape across different species subjected to different lifespan-extending interventions is represented by a fiber bundle, wherein individual degeneracy manifolds are continuous lines in the space of initial mortality rate *M*_0_ and mortality doubling rate *α*. The functional form of any given degeneracy manifold is well approximated by the Strehler-Mildvan dependence (Strehler and Mildvan, 1960), thus allowing us to identify the physical origin of parameters entering the Strehler-Mildvan correlation. We found that the Strehler-Mildvan dependence is one-rather than two-parametric, and that Strehler-Mildvan degeneracy manifolds for different species asymptotically approach each other in the limit of very slow aging rate *α* → 0.

We further determined that any given degeneracy manifold within the longevity landscape fiber bundle also represents a “sloppy” direction (Machta et al., 2013) in the space of parameters (*M*_0_, *α*); even moderate perturbations of lifespan distributions along such directions lead to a strong displacement in the space of survival curve parameters (*M*_0_, *α*). We found that the effects of most lifespan-modulating interventions correspond to the displacement along such degeneracy manifolds on the longevity landscape. This is consistent with recent observations that most interventions in large-scale screens do not affect the mortality doubling rate (the rate of aging) (Liu and Acar, 2018; Nowak et al., 2018; Pedro de Magalhães et al., 2018).

We further discovered that, for interventions leading to displacements along a given degeneracy manifold, there is a natural soft bound on the maximal achievable lifespan. We estimated the value of this bound for common model organisms and humans. The human bound (∼138 years) is higher than that recently reported (Brown et al., 2017; De Beer et al., 2017; Dong et al., 2016b; Hughes and Hekimi, 2017; Rozing et al., 2017) based on the data for the oldest old. It should be noted that our results for the bound are more general, as they follow from a statistical analysis of survival functions of entire cohorts rather than on the analysis of outliers. However, the bound is soft and may possibly be extended by interventions leading to a displacement of mortality curves away from the Strehler-Mildvan manifold characterizing a given species, such as in the case of calorie restriction in mice.

The physical origin of this bound is based on the finding that the mortality doubling rate *α* cannot be decreased without limit along the degeneracy manifold; at sufficiently small *α* the Kullback-Leibler divergence between the controlled and the perturbed distribution starts to increase providing a “soft bound” on the minimal achievable “rate of aging” *α*. These observations help explain why large-scale longevity screens produce few extremely long-lived outliers - in such screens, parameters of mortality curves are located along a single degeneracy fiber, and the maximal lifespan which can be achieved along a single such fiber is limited.

Interpretation of our estimated bound on human lifespan should be considered in the context of mortality data used to calibrate our model. Longevity data from model organisms are largely composed of single-gene (often null alleles) or single-parameter environmental/dietary variations measured under highly-controlled laboratory conditions in strains that have been inbred (often to homozygosity) and adapted to lab conditions for many generations. In contrast, the human population consists of individuals subjected to complex and dynamically changing environment. Individual genetic differences in humans also include many natural, non-lethal gene variants, but rarely homozygous null mutations for any given gene. Thus, the structure of Strehler-Mildvan degeneracy derived from human data reflects a different sampling of the “longevity space” compared to model organisms. One consequence of this is that, while many more genetic and environmental variants have been sampled in humans, the fraction of each variant in the population is relatively low. In contrast, many hundreds or even thousands of individual (sometimes genetically identical) yeast, worms, and mice were experimentally subjected to specific dietary regimens. Cohort sizes in those studies provided a sufficient statistical power to estimate precision of lifespan modulations by interventions.

Another important consideration is that the interventions giving the largest magnitude of lifespan extension in model organisms are almost certainly not sampled in the human population due to their substantial fitness costs. Consider, for example, Ames dwarf mice, which are among the longest-lived representatives of these species. They are unlikely to survive to the reproductive age in the natural environment due to defects in growth, development, and thermoregulation. Likewise, the *age-1* null worms that live ∼10 times longer than wild type *C. elegans* are sterile and fail to reach true adulthood. Even if such a mutant arose in the natural population, it would be rapidly selected out. Interestingly, these examples correspond to a displacement of parameters of mortality curves away from the main Strehler-Mildvan degeneracy manifold for the species. Although we are not aware of interventions which would lead to a similar displacement in humans, we believe that there may be extreme genetic and environmental interventions, perhaps akin to those sampled in model organisms, that could substantially exceed this bound. In particular, it may be expected that radical rejuvenation or biological engineering technologies will lead to this class of interventions. Thus, we interpret the bound of ∼138 years as a minimum estimate for maximal human longevity.

## Experimental Procedures

### Survival data for different species and interventions affecting lifespan

All demographic datasets used in the present study (with the exception of the screen for single gene knockdowns affecting lifespan of *D. melanogaster*, see below) were collected from publicly available sources. Those included: (i) survival data for 4,698 *S. cerevisiae* strains with single gene knockouts affecting replicative lifespan in yeast,(McCormick et al., 2015) (ii) survival data for *C. elegans* subjected to 1,416 different diets including individual pharmacological compounds from the LOPAC pharmacological network screen,(Ye et al., 2014) (iii) human mortality data from U.S. Social Security Administration Actuarial Tables and Human Mortality Database(Human Mortality Database, 2015), (iv) mortality and survival data for species presented in the COMADRE database(Salguero-Gómez et al., 2016) and (v) survival data from NIH Interventions Testing Program studying treatments with the potential to extend lifespan and delay disease in mice(Miller et al., 2007; Nadon et al., 2017; Strong et al., 2008; Warner et al., 2000). We also collected and aggregated mortality and survival data from different experiments on lifespan modulation in yeast, nematodes, fruit flies, mice and rats from literature (Table S1).

### Fitting survival data to Gompertz distributions

All fits were performed in Matlab using *fitlm* command with ‘RobustOpts’ option used to iteratively exclude outliers (typically, corresponding to deviations from Gompertz law at early and very late ages). Where possible, the fit errors were estimated by bootstrapping: randomly dividing cohort of animals into sub-cohorts, identifying parameters of Gompertz fit for the sub-cohorts and subsequently estimating mean and standard deviation of Gompertz parameters across sub-cohorts.

### Fitting Gompertz parameters to Strehler-Mildvan degeneracy manifold

All fits were performed in Matlab using *fitlm* command with ‘RobustOpts’ option used to iteratively exclude outliers. Outliers are reported in Figures in the main text and Supplementary Information.

### Age-dependent survival and mortality data from the COMADRE database

Survival and mortality data for species included in the COMADRE database(Salguero-Gómez et al., 2016) were determined using Caswell matrix population models.(Caswell, 2001; Keyfitz and Caswell, 2005) For every species included, the COMADRE database reports population model matrices *U* and *A*, the former encoding transitions and survival of extant individuals, while the latter, projection matrix defined in such a way that the population size at time *t* + 1 is related to the population size at time *t* according to the relation

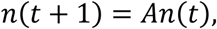

also including the contribution from production of individuals by clonal and sexual reproduction. The age-dependent survival and mortality rates were extracted directly from the matrix *U* (its diagonal elements giving the probability for individual specimen to transfer from the sub-cohort with age *a* to the sub-cohort with age *a* + 1) as well as from the eigenvector of the matrix *A* corresponding to a stable (or nearly stable) population dynamics *n*(*t*) ≈ *Const* (the structure of this eigenvector vector determines the lifespan distribution function of the cohort(Caswell, 2001)).

### Dietary perturbations for modulating lifespan of *Drosophila*

The lifespan screen for 162 single gene knockdowns modulating lifespan of *D. melanogaster* was performed in our lab. *Drosophila* species were obtained from UC San Diego Stock Center (La Jolla, CA, USA). The flies were maintained on corn meal food (85.7 g corn meal ‘Aunt Jemina’ (The Quaker Oats Company, Chicago, IL, USA), 50 ml golden A unsulfured molasses (Groeb Farms Inc, Onsted, MI, USA), 71.4 g Torula yeast (MP Biomedicals, Solon, OH, USA), 2.86 gp-hydroxybenzoic acid methyl ester(Sigma), 6.4 g agar (MoorAgar Inc, Loomis, CA, USA) and 5.7 ml propionic acid (Sigma) per liter of water). For lifespan experiments, approximately 50 young females of each strain/species were allocated per vial and 3 replicates were used per group. A total of two diet groups were used for lifespan determination of *Drosophila* species and strains. The first is basal and calorie restricted yeast/sugar based diet utilized frequently in fly lifespan experiments (Mair et al., 2005). The second is basal and amino acid restricted chemically defined diet (Lee et al., 2014).

### Genetic perturbations modulating lifespan of *Drosophila*

NAi studies were carried using GeneSwitch (GS) Gal4 System (Huang et al., 2014; Nicholson et al., 2008; Roman et al., 2001) and transgenic shRNA lines from Transgenic RNAi Resource Project (TRiP). The actin-GS-Gal4 virgin females were crossed to respective transgenic shRNA. The F1 progeny was raised at 25°C for 3-5 days to reach adulthood. Progeny were collected and separated according to genotype and gender with each assigned as control and RU486 (150 ug/mL). Adult flies were seeded at approximately 50 flies/vial, and 3 replicates were used per group. For lifespan experiments, flies were maintained on defined amino acid diets (Lee et al., 2014) with or without addition of RU486 at described concentration. Survivorship was recorded during food changes every 3 days when flies were transferred to new vials. Experiments were performed on both sexes. All species and strains were maintained in a temperature-controlled incubator at 25 °C with 12-h light/dark cycle and ∼60% humidity.

## Acknowledgments

The authors would like to thank Richard Miller for discussion and insightful comments. Supported by NIH AG047745.

## Figure legends

**Table 1.** Estimated maximal lifespans of humans and model organisms achievable along their associated Strehler-Mildvan degeneracy manifolds.

| Species | Maximal achievable lifespan |
| --- | --- |
| Budding yeast (replicative) | 147 generations |
| Nematode | 167 days |
| Fruit fly | 112 days |
| Mouse | 1480 days |
| Human | 138 years |

