## Supplementary File 1 for "The landscape of longevity across phylogeny"

**Supplemental File 1**

### 1. Kullback-Leibler divergence of Gompertz lifespan distribution, degeneracy on the fitting landscape and Strehler-Mildvan correlation

The nature of Strehler-Mildvan (SM) correlation was recently discussed in several publications (Burger and Missov, 2016; Tarkhov et al., 2017), including those suggesting that it is related to the existence of degenerate sub-manifolds in the space of parameters of Gompertz distribution. If so, SM correlation would have to be considered a fitting artifact and could not be used for quantitative population or evolutionary analysis of longevity. Here, we address the question of the existence of degenerate submanifolds in the space of Gompertz fits using standard tools of information-theoretical analysis.

To reiterate the argument on the nature of SM relationship (Tarkhov et al., 2017), assume that there is a number of data points $\{n_1, n_2, n_3, \ldots\}$ representing lifespans of individual organisms that belong to the same species. Mortality of the species is subject to Gompertz law $M(t) = -\frac{\frac{dS(t)}{dt}}{S} = M_0 \exp(\alpha t)$, and the lifespan distribution function is given by

$$F(t) = M_0 e^{\alpha t} \exp\left(\frac{M_0}{\alpha}(1 - e^{\alpha t})\right). \qquad (S1)$$

Assuming further that the lifespans $\{n_1, n_2, n_3, \ldots\}$ are produced from the distribution (S1) with particular values of parameters $x = \{M_0, \alpha\}$, one would now pick subsets such as $\{n_1, n_5, n_6, \ldots\}, \{n_2, n_3, n_7, \ldots\}$etc. from the dataset $\{n_1, n_2, n_3, \ldots\}$ and attempt to recover the values of Gompertz fitting parameters $\tilde{x} = \{\widetilde{M}_0, \tilde{\alpha}\}$ from the subsets. Apparently, the subsets $\{n_1, n_5, n_6, \ldots\}, \{n_2, n_3, n_7, \ldots\}$ etc. are produced from the very same lifespan distribution function, nevertheless according to (Tarkhov et al., 2017) one would be finding different values of $\{\widetilde{M}_0, \tilde{\alpha}\}$, although the values of $\widetilde{M}_0$ would be correlated with the values of $\tilde{\alpha}$.

Let us now determine the functional form of this correlation. Consider two distributions $F_1(t)$ and $F_2(t)$ characterized by the different Gompertz parameters $x = \{M_0, \alpha\}$. The quantitative question how much it is more likely for the set $\{n_1, n_2, n_3, \ldots\}$ to be described by the distribution $F_1(t)$ rather than the distribution $F_2(t)$ is addressed by estimating the Kullback-Leibler divergence (Kullback and Leibler, 1951) between the distributions $F_1(t)$ and $F_1(t)$

$$D_{KL}(F_1||F_2) = \int_0^{+\infty} dt \cdot F_1(t) \log\left(\frac{F_1(t)}{F_2(t)}\right) \qquad (S2)$$

For the Gompertz lifespan distribution (S1), the integral (S2) can be taken explicitly revealing:

$$D_{KL}(F_1||F_2) = \log\left(\frac{M_1}{M_2}\right) + e^{\frac{M_1}{\alpha_1}}\Gamma\left(0, \frac{M_1}{\alpha_1}\right)\left(1 - \frac{\alpha_2}{\alpha_1}\right) +$$

$$+\frac{M_2}{\alpha_2}\left(e^{\frac{M_1}{\alpha_1}}\left(\frac{\alpha_1}{M_1}\right)^{\frac{\alpha_2}{\alpha_1}}\Gamma\left(1 + \frac{\alpha_2}{\alpha_1}, \frac{M_1}{\alpha_1}\right) - 1\right) - 1 \qquad (S3)$$

where $\Gamma(a, x)$ is the incomplete Gamma function.

The measure $D_{KL}(F_1||F_2)$ is not symmetric with respect to the exchange $F_{1\to 2}(t)$ and cannot thus serve as a metric distance on the manifold of fitting parameters. For the latter purpose, it is more convenient to use the Fisher information matrix, which is obtained from $D_{KL}(F_1||F_2)$ by assuming that $F_1(t)$ and $F_2(t)$ are almost similar, with slightly different $\{M_0, \alpha\}$ and $\{M_0 + \delta M_0, \alpha + \delta\alpha\}$; one then finds

$$D_{KL}(F_1||F_2) \approx \frac{1}{2}\sum_{i,k} g_{ik}\delta x^i \delta x^k + O(\delta x^3),$$

where

$$g_{ik} = -\int_0^\infty dt \cdot F(t) \frac{\partial}{\partial x_i}\frac{\partial}{\partial x_j} \log(F(t)/\Lambda) \qquad (S4)$$

is the Fisher information matrix of the lifespan distribution density $F(t)$ and $x = \{M_0, \alpha\}$. Note that it is necessary to introduce a free parameter $\Lambda$, since the lifespan distribution density is a dimensionful quantity; however, all subsequent results are independent of this parameter.

The nature of possibly existing degenerate manifolds in the space of all Gompertz fits now becomes clear. If the Fisher information matrix (S10) has an eigenvalue very close to 0, all lifespan distributions obtained by translation along the corresponding direction in the space of parameters are practically identical per measures based on the Kullback-Leibler divergence.

We shall now determine the eigenvalues of the Fisher information matrix of the Gompertz lifespan distributions: $g_{ik} = \lambda_1 u_1 u_1^T + \lambda_2 u_2 u_2^T$, where $u_i$ are vectors in the metric space $\{M_0, \alpha\}$. In the regime $x = \frac{M_0}{\alpha} \ll 1$ (which is realized for the majority of species and perturbations discussed in this paper) one finds directly expanding the expression (S3) up to the second order in $\delta x$ and $\delta\alpha$:

$$\delta D_{KL} \approx \frac{1}{2}\left(\frac{\delta\alpha}{\alpha}\right)^2 \left(\log^2 x + (\gamma - 1)^2 + \frac{\pi^2}{6} + 2(\gamma - 1)\cdot \log x\right) +$$

$$+(1 - \gamma - \log x)\frac{\delta\alpha}{\alpha}\frac{\delta x}{x} + \frac{1}{2}\left(\frac{\delta x}{x}\right)^2, \qquad (S5)$$

where $\gamma = 0.57721566\ldots$ is the Euler-Mascheroni constant. One of the eigenvalues of the corresponding Fisher information matrix logarithmically vanishes in the limit $x \ll 1$ as

$$\lambda_1 \approx \frac{\pi^2}{12\log^4 x},$$

while another one remains logarithmically large:

$$\lambda_2 \approx \log^2 x.$$

Therefore, its corresponding eigenvector defines the direction of degeneracy. Indeed, one can write approximately (keeping only leading terms in large $|\log x|$):

$$\delta D_{KL} \approx \frac{1}{2}\left((\gamma - 1 + \log x)\frac{\delta\alpha}{\alpha} - \frac{\delta x}{x}\right)^2,$$

so that $\delta D_{KL}$ vanishes if the expression in the round brackets equals to zero (this expression then defines the differential equation with its solution identifying the direction of degeneracy, see Section 2 below).

The eigenvalue $\lambda_1$ grows with $x$, reaches maximum at $x \sim 1$ and then starts to decrease again at $x > 1$. On the other hand, $\lambda_2$ decreases in the interval $0 < x < 1$, reaches minimum at $x \sim 1$ and grows at $x > 1$. The two eigenvalues never become of the same order, and the eigenvector corresponding to the second eigenvalue $\lambda_2$ represents a degenerate (quasi-degenerate when the first eigenvalue $\lambda_1$ is not very small) manifold in the space of Gompertz parameters (Figure 4D of the main text).

**2. Degeneracy manifolds in the space of Gompertz parameters**

The form of the tangential Strehler-Mildvan curve determined by the differential (S5), which identifies a single degeneracy manifold in the space of Gompertz parameters, can be read straightforwardly from the expression for the Kullback-Leibler divergence (S3): it roughly corresponds to the first two terms in the Kullback-Leibler divergence, while the third term describes deviations from it. Thus, one has for the degeneracy manifold:

$$\log\left(\frac{M_1}{M_2}\right) + e^{\frac{M_1}{\alpha_1}}\Gamma\left(0, \frac{M_1}{\alpha_1}\right)\left(1 - \frac{\alpha_2}{\alpha_1}\right) \approx 0, \qquad (S6)$$

where $\alpha_1$ and $M_1$ (parameters of "control" distribution) are considered fixed. Then, it can be immediately seen that the parameters of the Strehler-Mildvan correlation are related to the parameters $M_1$ and $\alpha_1$ of the control distribution according to the expressions

$$K = M_1 \exp\left(e^{\frac{M_1}{\alpha_1}}\Gamma\left(0, \frac{M_1}{\alpha_1}\right)\right), \qquad (S7)$$

$$B = \frac{\alpha_1 e^{-\frac{M_1}{\alpha_1}}}{\Gamma\left(0, \frac{M_1}{\alpha_1}\right)}. \qquad (S8)$$

The third term given by

$$\delta D_{KL} = \frac{M_2}{\alpha_2}\left(e^{\frac{M_1}{\alpha_1}}\left(\frac{\alpha_1}{M_1}\right)^{\frac{\alpha_2}{\alpha_1}}\Gamma\left(1 + \frac{\alpha_2}{\alpha_1}, \frac{M_1}{\alpha_1}\right) - 1\right) - 1 \qquad (S9)$$

is the only non-trivial contribution to the Kullback-Leibler divergence when the equation (S6) holds (i.e., for interventions leading to lifespan perturbations along a single Strehler-Mildvan fiber).

In the limit $x = \frac{M_1}{\alpha_1} \gg 1$ (which corresponds to the case of long lifespans), one finds approximately

$$\delta D_{KL} \approx \left(\frac{M_2}{M_1} - 1\right)^2, \quad \frac{M_1}{\alpha_1} \gg 1. \qquad (S10)$$

and the degeneracy manifold (corresponding to vanishing $\delta D_{KL}$) is given by the condition $M \approx Const$ (see Fig. 4c of the main text).

In the limit $x = \frac{M_1}{\alpha_1} \ll 1$, which is realized for virtually all species and interventions considered in this paper, the expressions above simplify considerably, and we shall now proceed to the discussion of this case.

**3. Strehler-Mildvan degeneracy. Regime $x = \frac{M_0}{\alpha} \ll 1$**

In this regime, it is interesting to consider the case of a small deviation from a control lifespan distribution: $M_2 \approx M_1 + \delta M$, $\alpha_2 \approx \alpha_1 + \delta\alpha$, $\delta M \ll M_1$, $\delta\alpha \ll \alpha_1$. The condition (S6) then can be rewritten as a differential equation:

$$\alpha \frac{dx}{d\alpha} \approx x(\gamma - 1 + \log x), \qquad (S12)$$

where $x = M_0/\alpha$, indices 1,2 are dropped for simplicity so that $\alpha = \alpha_1$, $M_0 = M_1$, and the incomplete gamma function $\Gamma(0, x)$ is expanded at small $x$ (i.e., $\frac{M_0}{\alpha} \ll 1$, the regime which is realized for almost all species and interventions considered in this paper). Its solution can be found straightforwardly:

$$\log M_0 \approx \log\alpha - \frac{\alpha}{\tilde{B}} + 1 - \gamma \quad (S13)$$

or $\log x \approx -\frac{\alpha}{\tilde{B}} + 1 - \gamma$, confirming that the Strehler-Mildvan correlation is truly a one-parametric dependence rather than the two-parametric one.

This expression can be understood as follows. Fixing a point $(\alpha, x)$ on the landscape of Gompertz parameters, we fix a control lifespan distribution function, which we would like to study deviations from. With every such point an ellipse in the space of parameters $(\alpha, x)$ is associated with axes given by the eigenvalues of the Fisher information matrix of the control Gompertz distribution. The major axis of this ellipse is determined by the smaller eigenvalue $\lambda_1$ of the Fisher information matrix, while the minor axis – by the larger eigenvalue $\lambda_2$. Gompertz distributions with parameters within this ellipse do not deviate noticeably from the control distribution in the sense of Kullback-Leibler divergence, and thus fluctuations of Gompertz parameters within this ellipse are easy to achieve. Let us denote a single change of Gompertz parameters within this ellipse as an individual step of adjusting parameters of lifespan distribution by a particular lifespan-modulating intervention. The end point of the step represents a new Gompertz distribution which, as was explained above, deviates negligibly from the control one in terms of Kullback-Leibler divergence. We can then ask whether it is possible to continue adjustment process starting from the end point of the first step in such a way that newly obtained lifespan distributions will deviate negligibly from the ones obtained during the previous step, etc. The answer to this question is positive, and the form of the manifold including any such sequence of step is given by the expression (S13).

### 4. Approximate temporal scaling of lifespan distributions

Let us now consider scaling transformations of hazard functions and survival curves discussed in (Stroustrup et al., 2016). As it has been argued previously in (Stroustrup et al., 2016), hazard functions of cohorts of *C. elegans* maintained at two different temperatures $T_0$ and $T_1$ are related according to

$$h_{T_0}(\lambda^{-1}t) \approx \lambda h_{T_1}(t). \quad (S14)$$

As we have argued earlier, most perturbations of environment and genotype lead to displacement of parameters of the survival distribution function along the degeneracy lines corresponding to the condition $D_{KL}(F_1||F_2) = 0$ of vanishing Kullback-Leibler divergence between the distributions (which in turn reduces to the Strehler-Mildvan dependence in the limit $x = \frac{M_1}{\alpha_1} \ll 1$, see above). Naturally, we expect that small perturbations of the environment temperature belong to this class of perturbations.

By construction, the expression (S13) is equivalent to the Strehler-Mildvan degeneracy line (S6). Therefore, a perturbation of the lifespan distribution function along it, which corresponds to a scaling transformation $\alpha \to \lambda\alpha$, leads to a new lifespan distribution function with parameter $M_0{}'$ determined by $\log M_0' \approx \log(\lambda\alpha) - \frac{\lambda\alpha}{\tilde{B}} - \gamma + 1$. The corresponding transformation of the hazard function is

$$h(t) = M_0 e^{\alpha t} \to M_0' e^{\lambda\alpha t} = \lambda\alpha \cdot e^{-\frac{\lambda\alpha}{\tilde{B}} - \gamma + 1} e^{\lambda\alpha t}. \quad (S15)$$

Since the hazard function of the cohort maintained at a temperature $T_0$ at time rescaled according to $t \to \lambda^{-1}t$ is given by $h_{T_0}(\lambda^{-1}t) = \lambda\alpha \cdot e^{-\frac{\lambda\alpha}{\tilde{B}} - \gamma + 1} e^{\alpha t}$, we see that it coincides with the rescaled hazard function of the cohort maintained at the temperature $T_1$: $\lambda h_{T_1}(t) = \lambda\alpha \cdot e^{-\frac{\alpha}{\tilde{B}} - \gamma + 1} e^{\alpha t}$ up to the factor $e^{-(\lambda-1)\alpha/\tilde{B}}$, introducing *an additional temporal shift*. We conclude that in the regime $\frac{M_0}{\alpha} \ll 1$ of applicability of the Strehler-Mildvan relation perturbations corresponding to a displacement of parameters of lifespan distributions along degeneracy manifolds generally lead to a rescaling of these distributions of exactly the form reported in (Stroustrup et al., 2016) accompanied by an additional temporal shift $t \to t + (\lambda - 1)/\tilde{B}$.

### 5. Strehler-Mildvan degeneracy. More on the regime $\frac{M_0}{\alpha} \gg 1$

In the regime $\frac{M_0}{\alpha} \gg 1$ the Fisher information matrix reduces to the expression $\delta D_{KL} \approx \frac{1}{2}\left(\left(1 + \frac{1}{x}\right)\frac{\delta\alpha}{\alpha} + \frac{\delta x}{x}\right)^2$, vanishing if the differential equation

$$\frac{dx}{d\alpha} \approx -\frac{x+1}{\alpha}$$

is satisfied. Its general solution is given by

$$M_0 \approx C - \alpha.$$

Therefore, as was explained above, in the regime $x = \frac{M_0}{\alpha} \gg 1$ the degeneracy manifold is determined by the condition of constant initial mortality $M_0 \approx C$. The corresponding eigenvalue of the Fisher information matrix (S4)

$$\tilde{\lambda}_1 \approx \frac{1}{16x^2}$$

remains much smaller than the other one, which asymptotically approaches 1 at $x \to \infty$:

$$\tilde{\lambda}_2 \approx 1 + \frac{1}{x}.$$

Again, as two eigenvalues of the Fisher information matrix never become of the same order of magnitude, the eigenvalue $\lambda_1$ which remains small in the regime $x \ll 1$, matches to the eigenvalue $\tilde{\lambda}_1$ vanishing in the regime $x \gg 1$.

**6. A note on the existence of degeneracy manifolds corresponding to exact temporal scaling**

Here we would like to show that there exists no such $x = \frac{M_0}{\alpha}$ that *exact* scaling transformation corresponds to a degeneracy line on the landscape of longevity. Indeed, infinitesimal exact scaling is a transformation of the form $M_2 = (1 + \delta\lambda)M_1,\ \alpha_2 = (1 + \delta\lambda)\alpha_1$. Expanding expression (S3) to the second order in $\delta\lambda$ at arbitrary values of $x$ one finds:

$$\delta D_{KL} \approx \left(\frac{1}{2} + e^x \cdot G_{3,4}^{4,0}\left(x\middle|\begin{matrix}1,1,1\\0,0,0,2\end{matrix}\right)\right)\delta\lambda^2 + O\big((\delta\lambda^3)\big),$$

where $G$ is Mejier G-function (Abramowitz and Stegun, 1972). It can be immediately checked by numerical simulation that the factor in front of $\delta\lambda^2$ is positive (and larger than $\frac{1}{2}$ for arbitrary $x$) implying that scaling never corresponds to an exact flat direction on the landscape of longevity. However, when $x \gg 1$ (the case of relatively short-lived species) one approximately has $\delta D_{KL} \approx \frac{1}{2}(\delta\lambda)^2$, meaning that scaling transformation indeed becomes nearly degenerate with a small effect on the value of Kullback-Leibler divergence between the two lifespan distributions (related by a scaling transformation) unless the scale factor becomes particularly large.

**7. The bounds on the maximal and average lifespans achievable along a single Strehler-Mildvan degeneracy manifold**

Generally, Gompertz lifespan distributions located along the same degeneracy manifold (S6) correspond to slightly different cases of population dynamics, simply because the mean and maximal lifespans calculated using those distributions differ. Indeed, recall that the mean lifespan derived using the Gompertz lifespan distribution (S1) is given by

$$E(t_{LS}) = \frac{1}{\alpha} e^{\frac{M_0}{\alpha}} \Gamma\left(0, \frac{M_0}{\alpha}\right), \qquad (S16)$$

clearly varying with translations along (S6). For the median lifespan, one finds instead a simpler expression

$$Median(t_{LS}) = \frac{1}{\alpha}\log(1 + \frac{\alpha}{M_0}\log 2).$$

The maximal lifespan $LS_{max}$ can be estimated fixing the size of cohort $N_0$; it is then given by the requirement that only a single organism from the cohort would survive by the time $\geq LS_{max}$. Performing the calculation and using the Strehler-Mildvan correlation, one then finds that

$$LS_{max} \approx \frac{1}{\alpha}\log(1 + \frac{\alpha}{K}e^{\alpha/B}\log(N_0)). \qquad (S17)$$

This expression also obviously varies, when subjected to translations along (S6). In particular, since both expressions (S16) and (S17) are monotonic functions of $\alpha$, both average and maximal lifespans monotonically grow along the Strehler-Mildvan fiber when the value of $\alpha$ is decreased.

Note, however, that for Gompertz lifespan distributions located along the degeneracy manifold (S6) are characterized by slightly different values of Kullback-Leibler divergence from the control Gompertz distribution, which fixes the position of the degeneracy manifold. This slight difference is determined by the term (S9) in the full Kullback-Leibler divergence: for translations along the Strehler-Mildvan degeneracy manifold/fiber we find

$$\delta D_{KL} \approx \frac{K}{B}\exp\left(-\frac{\alpha_2}{B}\right). \qquad (S18)$$

As one can see, this contribution to the Kullback-Leibler divergence grows with decreasing $\alpha_2$, the degeneracy manifold ceases to be "sloppy" at sufficiently small $\alpha_2$. In other words, Gompertz lifespan distributions deviating by more than $\delta D_{KL} \sim 1$ from the control distribution will not be easily achieved by perturbations displacing parameters of lifespan distributions along the same Strehler-Mildvan degeneracy manifold/ fiber. Thus, the associated minimal $\alpha$ achievable by translations along this fiber can be estimated as

$$\alpha_{min} \approx B \cdot \log\left(\frac{K}{B}\right), \qquad (S19)$$

and expressions for the average and maximal lifespans achievable by such translations read

$$\max\big(E(t_{LS})\big) \approx \frac{1}{B}, \qquad (S20)$$

$$\max(LS_{max}) \approx \frac{1}{B\log(K/B)}\log(1 + \log\left(\frac{K}{B}\right) \cdot \log(N_0)). \qquad (S21)$$

We note in passing that for all species considered in the present paper the condition $\frac{K}{B} > 1$ holds comfortably.

The bound (S21) is soft. First, it weakly (as a double logarithm) depends on the initial size of the cohort $N_0$ and will thus grow, albeit very slowly, with increased $N_0$. Second, it is based on the criterion $\delta D_{KL} \sim 1$, and higher values of $\delta D_{KL}$ cutoff can be used in principle, achieved by perturbations with potential to displace parameters of the survival curve from the current degeneracy line/fiber. Nevertheless, the value (S17) is robust as it depends on $N_0$ extremely weakly.

#### 8. Relevant expressions in the case $x \ll 1$

Most expressions derived above are simplified significantly in the limit $x = \frac{M_0}{\alpha} \ll 1$, which is realized for most species and interventions considered in the manuscript. Since $\log M_0/\alpha \approx -\frac{\alpha}{B} + 1 - \gamma$ on the Strehler-Mildvan degeneracy line in the limit $x \ll 1$, $|\alpha/B|$ is large ($\gg 1$) in the same limit. We thus find to the leading order in $\left|\frac{\alpha}{B}\right| \gg 1$ that the mean lifespan now reads

$$E(t_{LS}) \approx \frac{1}{B} - \frac{1}{\alpha}, \qquad (S21)$$

while the median lifespan is reduced to

$$Median(t_{LS}) \approx \frac{1}{B} + \frac{1}{\alpha}(\gamma - 1 + \log\log 2) \approx \frac{1}{B} - \frac{0.5772}{\alpha}. \qquad (S22)$$

Thus, estimating mean and median lifespans only from the lifespan distributions instead of the full set of Gompertz parameters approximately lifts the Strehler-Mildvan degeneracy (as was pointed out in (Tarkhov et al., 2017)), although degeneracy lifting is not complete with an additive strongly fluctuating contribution of the order of small parameter $B/\alpha$. The main contribution to both mean and median lifespans in this regime comes from the Strehler-Mildvan slope $\sim 1/B$. We also note that (a) in the limit $\alpha \to \infty$ median and mean lifespans coincide and are fully determined by the value of Strehler-Mildvan slope, (b) decreasing the rate of aging $\alpha$ leads to a *decrease* in both mean and median lifespans.

For the minimum aging rate $\alpha_{min}$ achievable by translations along the one-parametric Strehler-Mildvan degeneracy manifold (S13) we find

$$\alpha_{min} \approx (1 - \gamma)B \approx 0.4228 \cdot B, \qquad (S23)$$

roughly corresponding to the case of breakdown of the approximation $x \ll 1$ (note that at $\alpha = \alpha_{min}$ the expressions (S21), (S22) are inapplicable). We then obtain the following expression for the maximal lifespan achievable by translations along a particular Strehler-Mildvan manifold:

$$LS_{max} \approx \frac{1}{(1-\gamma)B} \log\left(1 + e^{\gamma - 1 + \frac{\alpha}{B}} \log N_0\right) \approx \frac{1}{(1-\gamma)B} \log\log N_0. \qquad (S24)$$

#### 9. Weak deviations from Gompertz lifespan distribution

Consider the case where the control lifespan distribution function $F_1(t) = f_1(t; M_0, \alpha)$ is again Gompertz-like but the new distribution to be compared to the control has the form $F_2(t) = f_2(t; M_0', \alpha') + \epsilon\delta f(t)$, where the contribution $\epsilon\delta f(t)$ is non-Gompertz-like but small, this smallness being regulated by the dimensionless parameter $\epsilon$. This case corresponds to a physical situation where the second distribution can still be sufficiently well approximated by the Gompertz one; the residuals of the corresponding Gompertz fit represented by the term $\epsilon\delta f(t)$ are small but not necessarily random. Note that $\int_0^{+\infty} dt\, \delta f(t) = 0$ by construction since both $f_1(t; M_0, \alpha)$ and $f_2(t; M_0', \alpha')$ are normalized to 1.

The Kullback-Leibler divergence between the two distributions $F_1(t)$ and $F_2(t)$ will thus have the form (to the first order in $\epsilon$)

$$D_{KL} = \delta D_{KL}(M_0, \alpha || M_0', \alpha') - \int_0^{+\infty} dt f_1(t; M_0, \alpha) \log\big(1 + \epsilon\delta f(t)\big), \quad (S25)$$

where $\delta D_{KL}(M_0, \alpha || M_0', \alpha')$ is given by the expression (S3). The second term in (S25) is the r.h.s. is the Kullback-Leibler divergence between the distributions $f_1(t; M_0, \alpha)$ and $f_1(t; M_0, \alpha)(1 + \epsilon\delta f(t))$, again properly normalized to 1 if the integral $\epsilon \int_0^{+\infty} dt f_1(t; M_0, \alpha)\delta f(t)$ vanishes (if this is not the case, the modification of the expression above is obtained by a subtraction of the constant term $\sim \log \int dt f_1(t)(1 + \epsilon\delta f(t))$ from the full Kullback-Leibler divergence (S25)). It is thus positive semidefinite at any $M_0$ and $\alpha$. On the other hand, the whole expression (S25) as well as the first term $\delta D_{KL}$ are also positive semidefinite. Therefore, if one looks for the degenerate manifolds defined by the condition $\delta D_{KL} = 0$ (vanishing of the first term in (S25)), the weak modification of the Gompertz distribution does not prohibit existence of degenerate manifolds or influence their locations in the space of parameters $(M_0, \alpha)$ much. However, it does generally influence the "widths" of degeneracy manifolds defined by the condition $D_{KL} \lesssim 1$ (Section 4 of these Supplementary Materials) and shifts the overall bias in $D_{KL}$, but this influence is perturbatively small in powers of $\epsilon$.

More accurately, we come to the same conclusion when considering $\epsilon$ as an additional parameter present in the distributions to be compared and estimating the Fisher information matrix by expanding the expression (S25) in small $\delta\alpha$, $\delta M_0$ and $\delta\epsilon$. If the integrals $\int_0^{+\infty} dt\, f_1 \delta f^2$, $\int_0^{+\infty} dt\, \frac{\partial f_1}{\partial \alpha} \delta f$ and $\int_0^{+\infty} dt\, \frac{\partial f_1}{\partial M_0} \delta f$ vanish or remain small, the only effect of residuals of the Gompertz fit on the Fisher information matrix is the emergence of a third eigenvalue, which also vanishes. In particular, this is the case if the weak modifications of Gompertz law are realized at late times $t > E(t_{LS})$ (where $E(t_{LS})$ is given by the expression (S16)). Unless the function $\delta f(t)$ grows very rapidly with age (in which case the perturbation $\delta f$ can naturally no longer be considered small), the three integrals above will remain bounded as the function $f_1(t)$ vanishes super-exponentially. In other words, in that case the functions $\delta f(t)$ and $\delta f^2(t)$ have a small support in the integrals $\int_0^{+\infty} dt\, f_1 \delta f^2$, $\int_0^{+\infty} dt\, \frac{\partial f_1}{\partial \alpha} \delta f$ and $\int_0^{+\infty} dt\, \frac{\partial f_1}{\partial M_0} \delta f$. The same applies to weak modifications of Gompertz law in the limited interval of early ages. To conclude, the formal smalless of these three integrals with respect to integrals entering the definition of Fisher information matrix for purely Gompertz lifespan distribution functions is the criterion of the smallness of perturbation $\delta f(t)$ which we use in this Section.

An important particular case of a large, rapidly growing perturbation $\delta f$ is when the mortality rate $M(t)$ corresponding to the lifespan distribution function $F_2(t)$ approaches plateau (so that the function $F_2(t)$ itself vanishes exponentially, rather than super-exponentially, at large $t$). As was discussed in the main text and Section 6 of Supplementary Materials, this case can be embedded in the developed framework as a non-Strehler-Mildvan sector on the longevity landscape, corresponding to $x = \frac{M_0}{\alpha} \gg 1$.

Finally, another argument in favor of insensitivity of Kullback-Leibler divergence-based analysis with respect to weak perturbations of Gompertz lifespan distributions is behavior of invariants of the Fisher information matrix under generic reparameterizations of lifespan distributions $(M_0, \alpha) \to (y_1, y_2)$ which encode generic changes in lifespan distributions. It is well known that under such reparameterizations scalar curvature constructed from the Fisher informatic matrix used as a metric tensor remains invariant. In the case of two parameters ($(M_0, \alpha)$ such as in our case) the scalar curvature remains a function of the determinant of the metric tensor only, i.e., the product of eigenvalues of Fisher information matrix, which vanishes at arbitrary $x$ as we have seen. Thus, general non-singular perturbations of Gompertz lifespan distribution functions which can be represented as reparameterizations $(M_0, \alpha) \to (y_1, y_2)$ will keep the scalar curvature of the Fisher information matrix vanishing implying existence of degeneracy manifolds in a general two-parametric fitting space of lifespan distribution functions. Again, a perturbation of Gompertz mortality curve leading to a plateau of mortality at late ages is an exception from this rule, since it belongs to a class of singular perturbations.

### Figure supplements

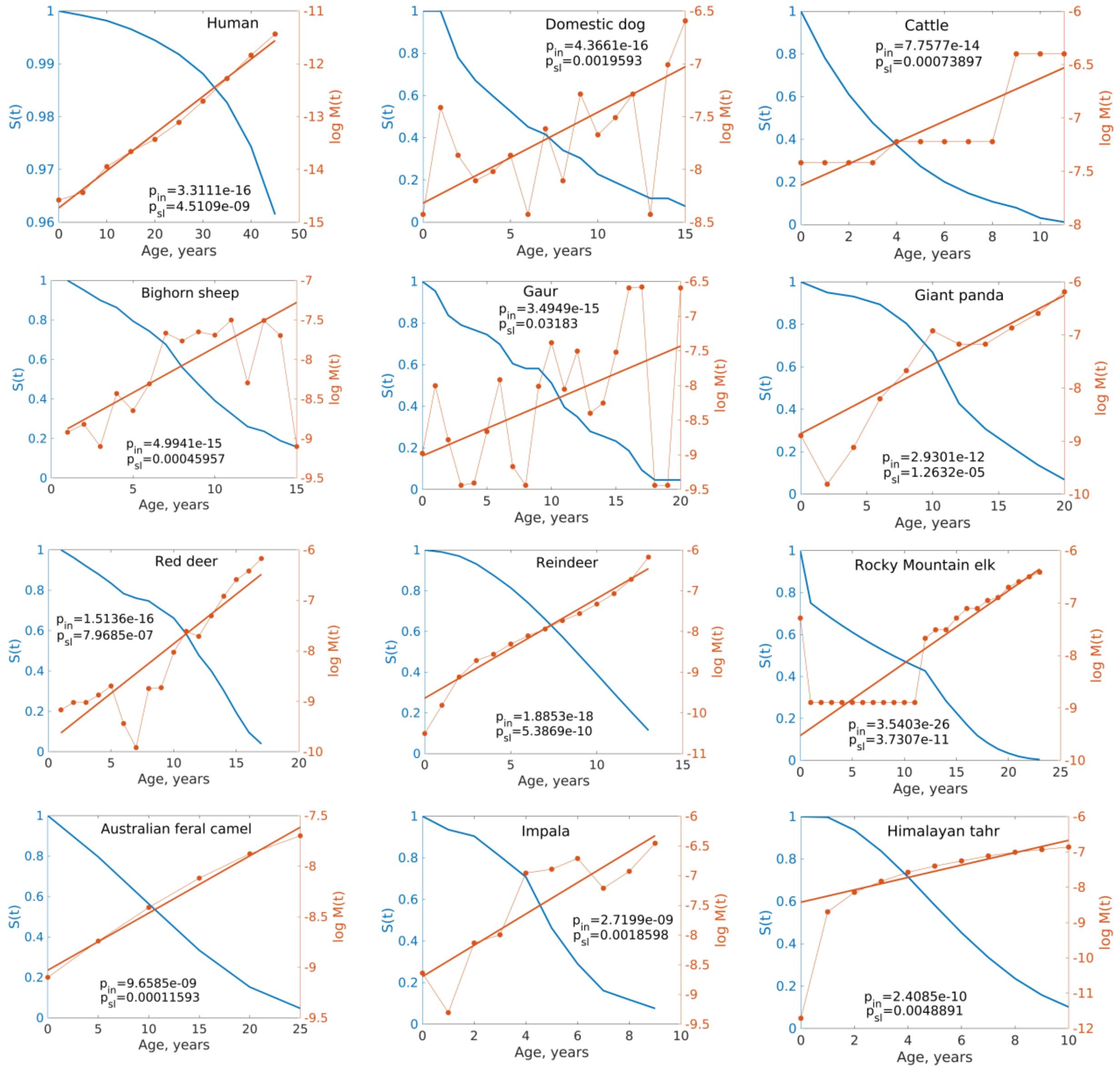


**Figure 1 – figure supplement 1.** Survival curves of various long-lived species reconstructed from COMADRE database. Blue curves – survival $S(t)$ as a function of age $t$, red dotted curves – natural logarithm of age-dependent mortality $M(t) = \log(-\frac{dS(t)}{dt}/S(t))$, red straight lines – robust linear regression $\log M(t) = \alpha_0 + \alpha_1 t + \epsilon$ of $\log M(t)$ to age, $p_{in}$ – p-value for the intercept coefficient $\alpha_0$ in the regression, $p_{sl}$ – p-value for the slope coefficient $\alpha_1$ in the regression.

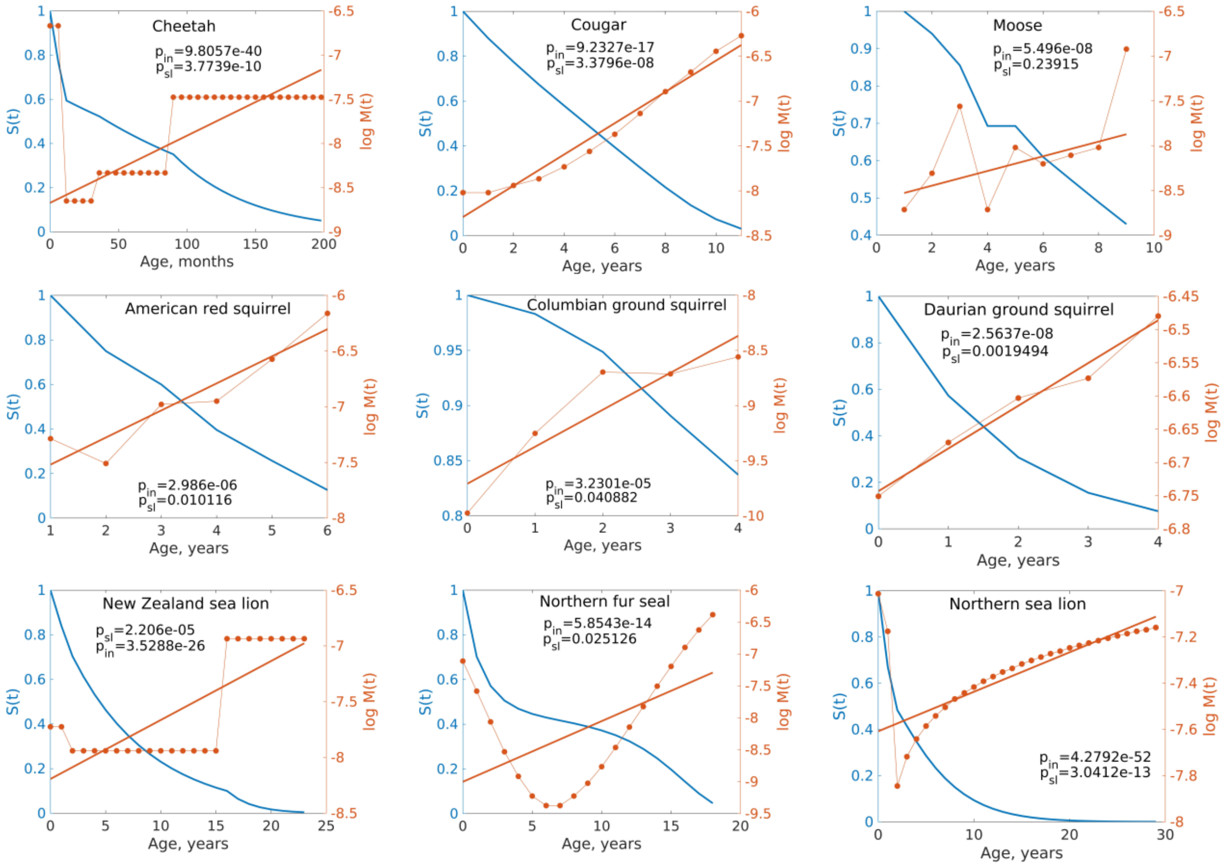


**Figure 1 – figure supplement 2.** Survival curves of various species with lifespans from several years to several decades reconstructed from COMADRE database. Blue curves – survival $S(t)$ as a function of age $t$, red dotted curves – natural logarithm of age-dependent mortality $M(t) = \log(-\frac{dS(t)}{dt}/S(t))$, red straight lines – robust linear regression $\log M(t) = \alpha_0 + \alpha_1 t + \epsilon$ of $\log M(t)$ to age, $p_{in}$ – p-value for the intercept coefficient $\alpha_0$ in the regression, $p_{sl}$ – p-value for the slope coefficient $\alpha_1$ in the regression.

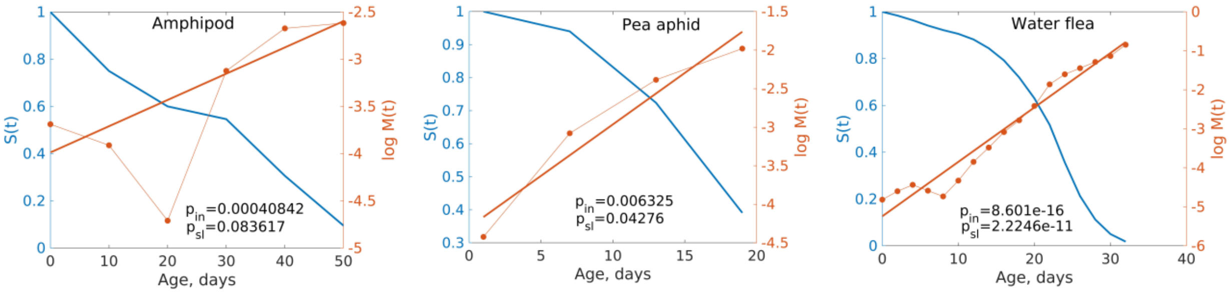


**Figure 1 – figure supplement 3.** Survival curves of several species with relatively short lifespans (~ several days) reconstructed from COMADRE database. Blue curves – survival $S(t)$ as a function of age $t$, red dotted curves – natural logarithm of age-dependent mortality $M(t) = \log(-\frac{dS(t)}{dt}/S(t))$, red straight lines – robust linear regression $\log M(t) = \alpha_0 + \alpha_1 t + \epsilon$ of $\log M(t)$ to age, $p_{in}$ – p-value for the intercept coefficient $\alpha_0$ in the regression, $p_{sl}$ – p-value for the slope coefficient $\alpha_1$ in the regression.

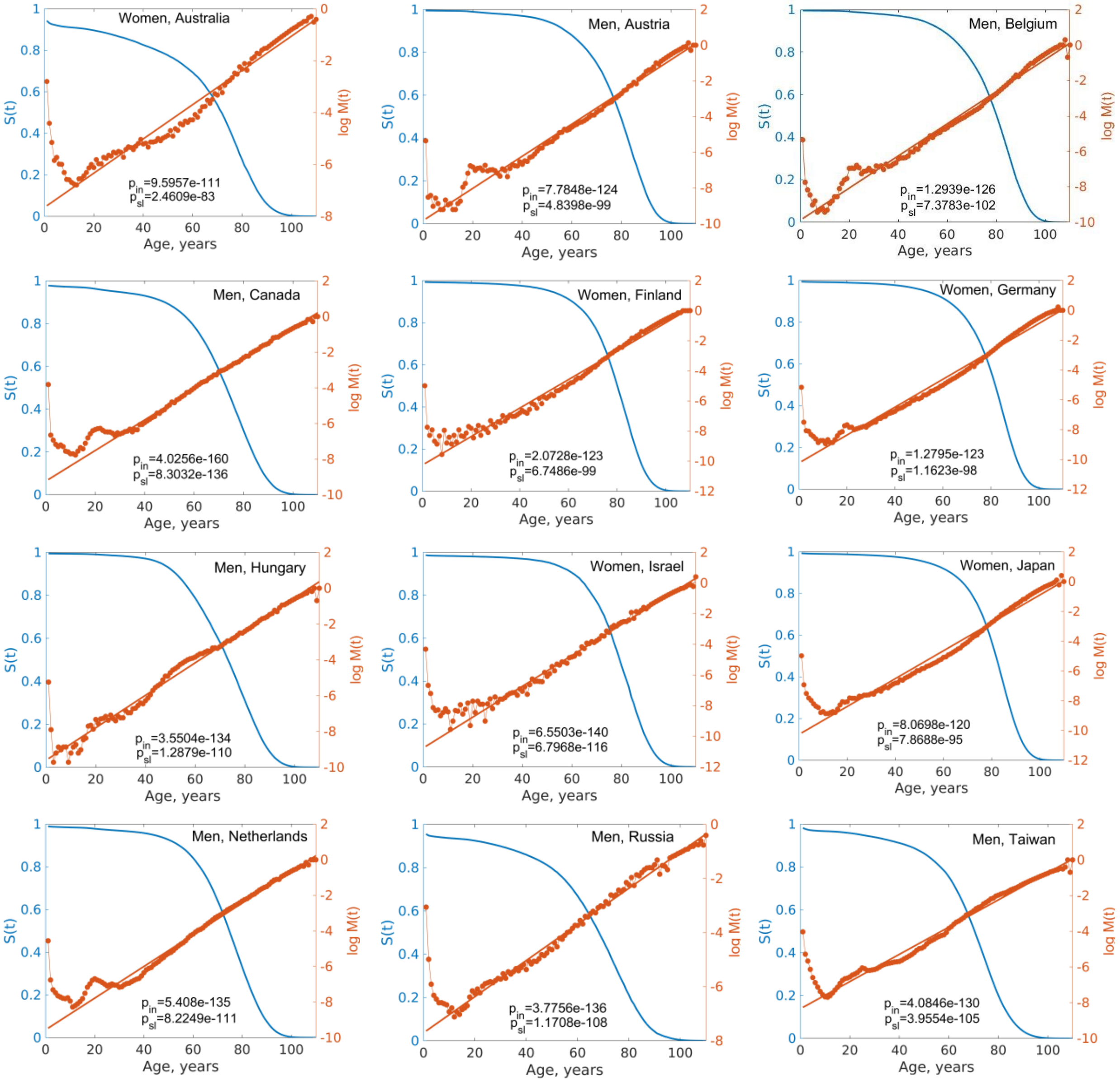


**Figure 1 – figure supplement 4.** Human survival curves (both men and women) estimated using demographic data collected in different countries (data from Human Mortality Database). Blue curves – survival $S(t)$ as a function of age $t$, red dotted curves – natural logarithm of age-dependent mortality $M(t) = \log(-\frac{dS(t)}{dt}/S(t))$, red straight lines – robust linear regression $\log M(t) = \alpha_0 + \alpha_1 t + \epsilon$ of $\log M(t)$ to age, $p_{in}$ – p-value for the intercept coefficient $\alpha_0$ in the regression, $p_{sl}$ – p-value for the slope coefficient $\alpha_1$ in the regression.

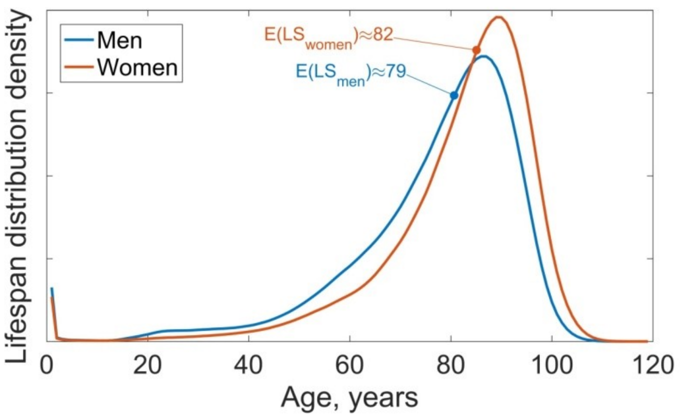


**Figure 1 – figure supplement 5.** Lifespan distribution function for men (blue) and women (red). Data from Social Security Administration actuarial tables.

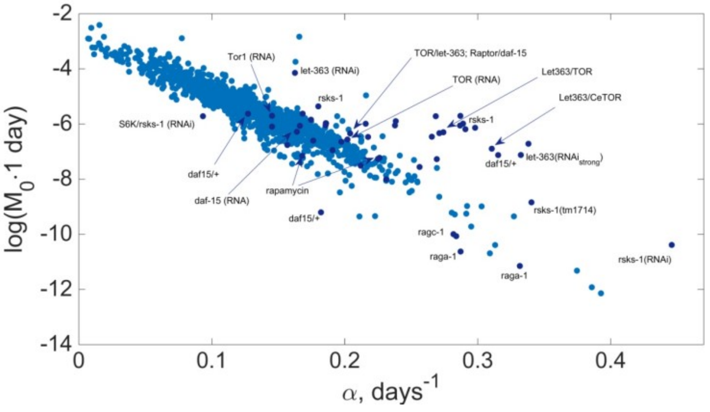


**Figure 2 – figure supplement 1.** Combining mortality curve parameters for *C. elegans* (Ye et al., 2014), shown in light blue, with the data from other published experiments (shown in dark blue, Table S1).

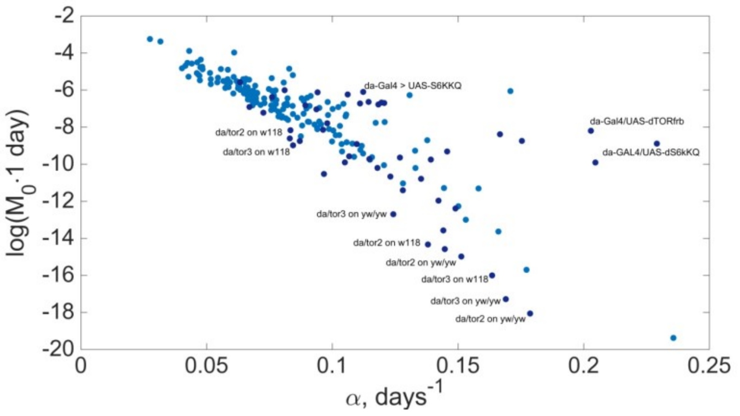


**Figure 2 – figure supplement 2.** Combining mortality curve parameters for fruit flies from our experiments (160 gene knockdowns), shown in light blue, with the data from other experiments (shown in dark blue, Table S1). Note that data points corresponding to *da/tor3* and *da/tor3* knockdowns on different backgrounds can be effectively combined into a separate SM line corresponding to lower $\log(M_0)$. Unmarked dark points correspond to rapamycin treatments with different concentrations.

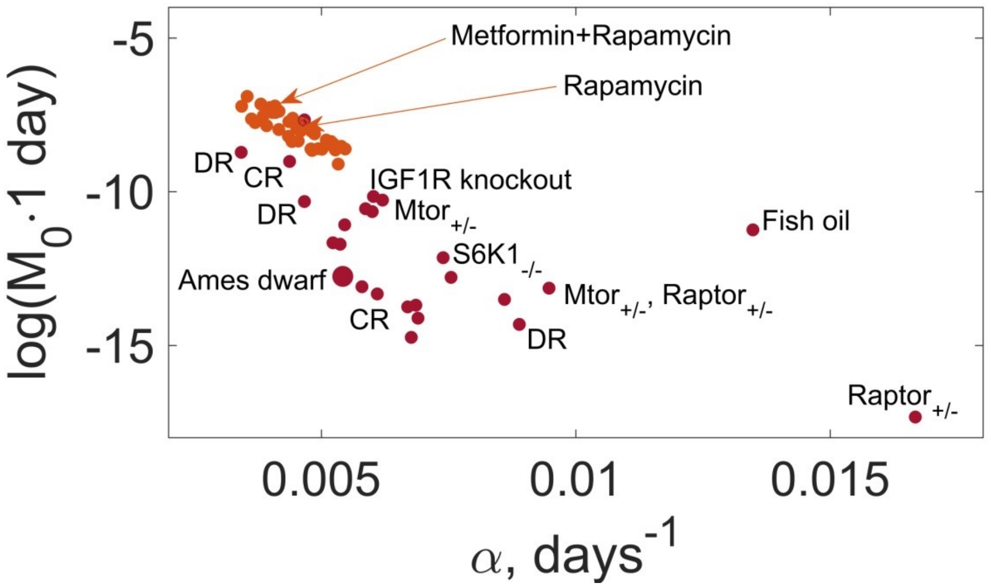


**Figure 2 – figure supplement 3.** Combining mortality curve parameters for mice (females) from the ITP program, shown in orange, with other data reported in the literature (shown in dark red, Table S1). The extreme long-lived Ames mice are shown with a larger dot. Most unmarked dark red dots correspond to various calorie restriction and dietary restriction experiments (note the displacement from the main SM line).

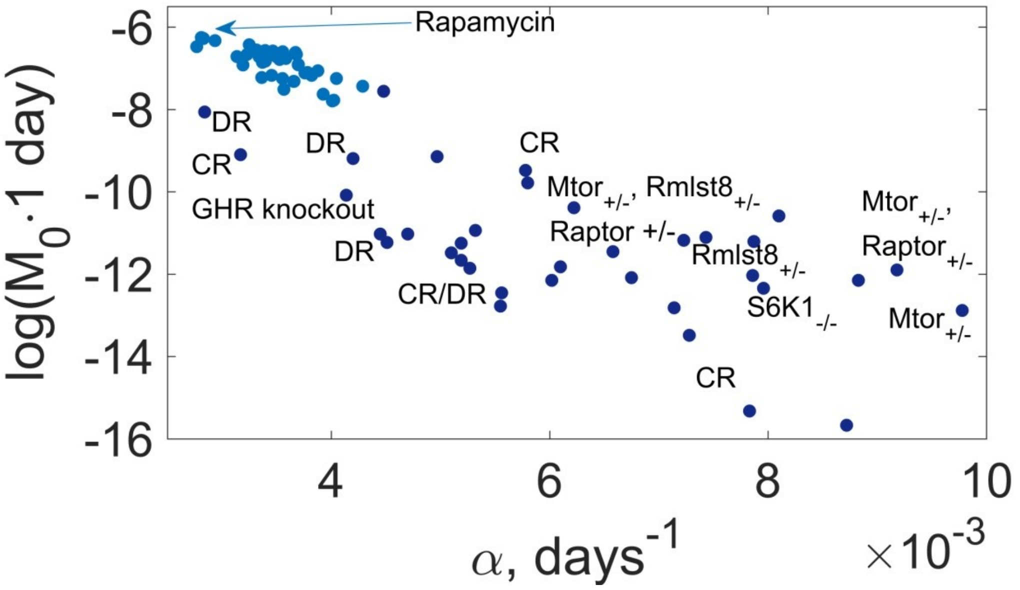


**Figure 2 – figure supplement 4.** Combining mortality curve parameters for mice (males) from the ITP program, shown in light blue, with other data reported in the literature (shown in dark blue, Table S1). Most unmarked dark blue dots correspond to various calorie restriction and dietary restriction experiments (note the displacement from the main SM line).

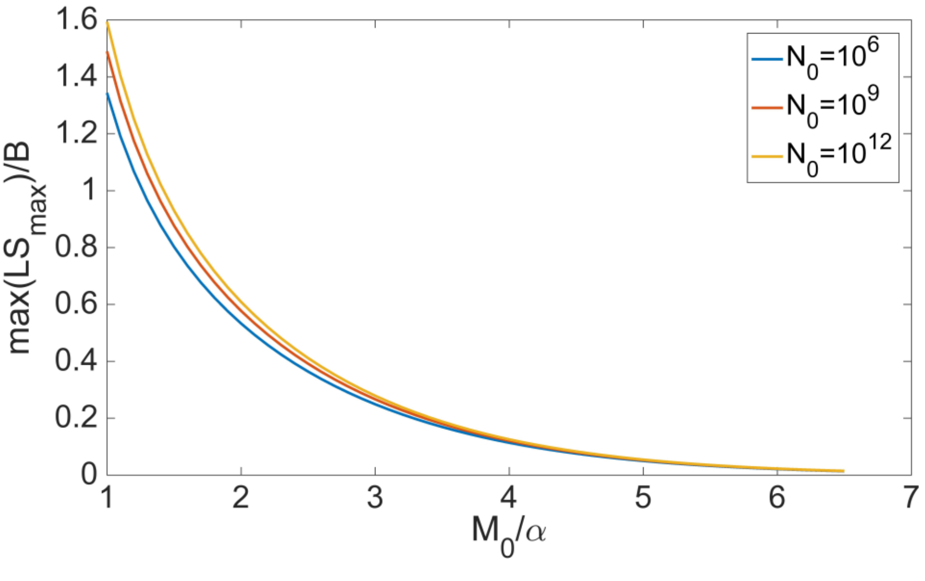


**Figure 3 – figure supplement 1.** Maximal value of $LS_{max}$ achievable along a given degeneracy manifold/fiber in units of the Strehler-Mildvan slope $B$ as a function of the dimensionless parameter $M_0/\alpha$ describing the control distribution; curves for several different initial cohort sizes are presented.

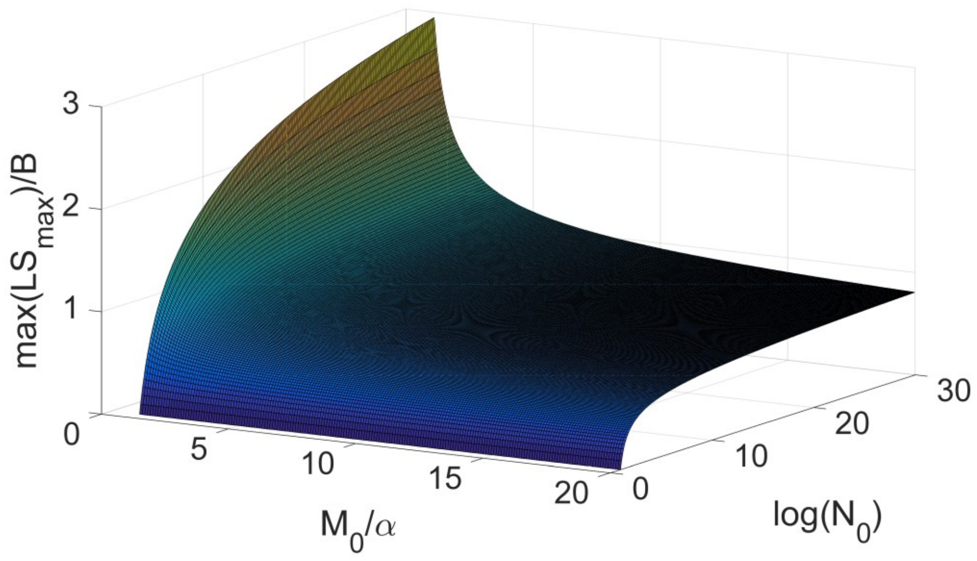


**Figure 3 – figure supplement 2.** The same as a function of $M_0/\alpha$ and the logarithm of the initial cohort size $\log(N_0)$.

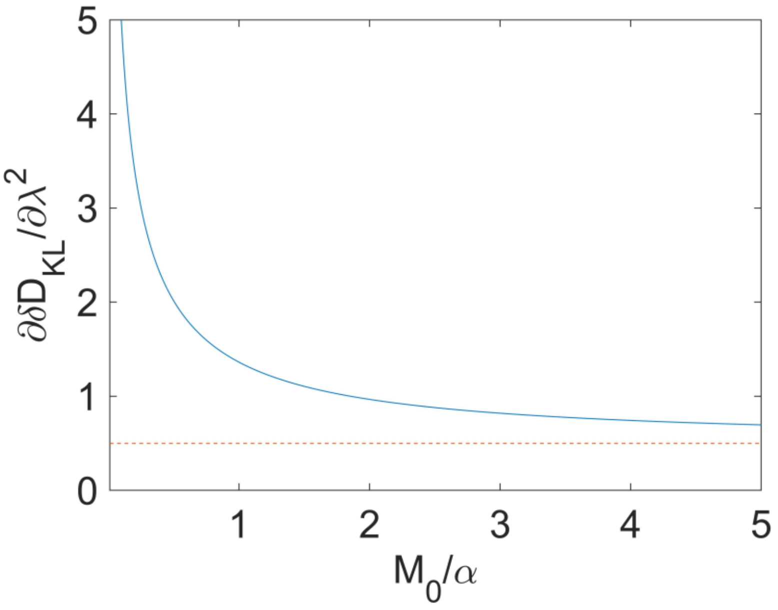


**Figure 3 – figure supplement 3.** Behavior of the Fisher information matrix for two Gompertz distributions related by a scaling distribution $M_0 \to \lambda M_0$, $\alpha \to \lambda\alpha$ as a function of $x = \frac{M_0}{\alpha}$.
